# LRTK: A platform agnostic toolkit for linked-read analysis of both human genomes and metagenomes

**DOI:** 10.1101/2022.08.10.503458

**Authors:** Chao Yang, Zhenmiao Zhang, Yufen Huang, Xuefeng Xie, Herui Liao, Jin Xiao, Werner Pieter Veldsman, Kejing Yin, Xiaodong Fang, Lu Zhang

## Abstract

Linked-read sequencing technologies generate high base quality reads that contain extrapolative information on long-range DNA connectedness. These advantages of linked-read technologies are well known and has been demonstrated in many human genomic and metagenomic studies. However, existing linked-read analysis pipelines (e.g., Long Ranger) were primarily developed to process sequencing data from the human genome and are not suited for analyzing metagenomic sequencing data. Moreover, linked-read analysis pipelines are typically limited to one specific sequencing platform. To address these limitations, we present the Linked-Read ToolKit (LRTK), a unified and versatile toolkit for platform agnostic processing of linked-read sequencing data from both human genomes and metagenomes. LRTK provides functions to perform linked-read simulation, barcode error correction, read cloud assembly, barcode-aware read alignment, reconstruction of long DNA fragments, taxonomic classification and quantification, as well as barcode-assisted genomic variant calling and phasing. LRTK has the ability to process multiple samples automatically, and provides the user with the option to generate reproducible reports during processing of raw sequencing data and at multiple checkpoints throughout downstream analysis. We applied LRTK on two benchmarking and three real linked-read data sets from both the human genome and metagenome. We showcase LRTK’s ability to generate comparative performance results from the preceding benchmark study and to report these results in publication-ready HTML document plots. LRTK provides comprehensive and flexible modules along with an easy-to-use Python-based workflow for processing linked-read sequencing datasets, thereby filling the current gap in the field caused by platform-centric genome-specific linked-read data analysis tools.

## Introduction

Linked-read sequencing generates reads with high base quality and extrapolative information on long-range DNA connectedness, which has led to significant advancements in human genome and metagenome research[1–3]. It circumvents the typical lack of long-range DNA information in short-read sequencing, and the high error rates and large initial DNA load requirements of long-read sequencing (e.g., Oxford Nanopore and Pacific Bioscience). These advantages of linked-read sequencing are invaluable when dealing with challenging cases of low-input clinical samples, such as cancer tissues or infectious disease samples. Linked-read technology furthermore promotes haplotype construction and the detection of complex structural variations[4], and its relatively low cost enables the application in large cohort studies.

Linked-read sequencing platforms, such as 10x Genomics linked-read (10x Genomics; now discontinued) and the newly developed single-tube long fragment read (stLFR)[2] and transposase enzyme linked long-read sequencing (TELL-Seq)[3], hold much promise in the metagenomics area. The hidden long-range information they provide enables local assembly of co-barcoded reads and thus significantly increases the number of high quality MAGs[5]. In longitudinal sequencing data sets[6–8], the barcodes associated with linked-read promote the phasing of variants and refine the identification of intra-host evolution of gut microbiota. In some complex environments, such as soil, linked-read sequencing has been shown to aid in the investigation of soil microbe genomes[9]. However, existing linked-read pipelines are mainly designed for use with the human genome, which points to an urgent need for appropriate metagenome analysis toolkits **(Table S1)**.

Despite the limitations that genome specificity places on research scope, linked-read sequencing has already been successfully applied to a myriad of genomic studies. For example, Long Ranger[10] performs barcode-aware read alignment and implements modules for variant calling and phasing using 10x Genomics linked-read. Tell-Sort[3] is a Docker-based pipeline to process raw TELL-Seq reads, and detect and phase genomic variants. stLFR[2] has found application in a customized pipeline that has been developed to first convert its raw reads into a 10x-compatible format, after which Long Ranger is applied for downstream analysis. The former pipeline however typically requires a lot of random-access memory and its data format conversion process is time consuming. Beyond the preceding examples that validate the inherent usefulness of linked-read technology, our search of the literature furthermore revealed a lack of unified and open-source toolkits that are compatible with the different linked-read platforms.

To this end, we present Linked-Read ToolKit (LRTK), a unified and versatile toolkit to analyze both metagenome and human genome linked-read sequencing data derived from any of the three major linked-read sequencing platforms. LRTK delivers a suite of utilities to simulate linked-read sequencing data, barcode correction, read cloud assembly, barcode-aware alignment, reconstruction of long fragments, and variant detection and phasing. LRTK is open-source, automatically produces HTML reports to summarize quality statistics as part of its pipeline, and generates publication-ready visualizations. We applied LRTK on benchmarking metagenome and human genome sequencing data (ATCC-MSA-1003 and NA12878) to evaluate the performance of 10x Genomics, stLFR and TELL-Seq linked-read sequencing technologies. We also applied LRTK on three real data sets to demonstrate its potential applications. Our results show that LRTK performs favorably on human genomic sequencing data when compared to single platform technologies, and that it adequately allows for data analysis of metagenomic sequencing data.

## Results

### Overview of LRTK

We developed a comprehensive workflow (which we refer to as LRTK) that takes raw linked reads from 10x Genomics, stLFR, or TELL-Seq technologies, and processes these inputs in a multi-step checkpointed pipeline that ends with the generation of user-friendly reports. LRTK consists of two main sections for metagenomic and human genome sequencing data (**Figure 1**). For metagenomic sequencing, LRTK includes the representative genomes from the UHGG[11] project as its default microbial reference genomes. LRTK uses a modified barcode-aware aligner based on EMA[12] to map reads to the reference genomes. To reduce mapping errors, LRTK employs a tiered alignment approach to align the sequencing reads. First, it identifies candidate genomes from the alignment files, and extracts the related genomes from the reference genome database. Second, the reads are aligned to the selected reference genomes to provide correct metagenomic read assignments. If the single nucleotide variant (SNV) calling function is enabled, LRTK also performs variation calling and phasing for the identified species.

**Figure 1.**
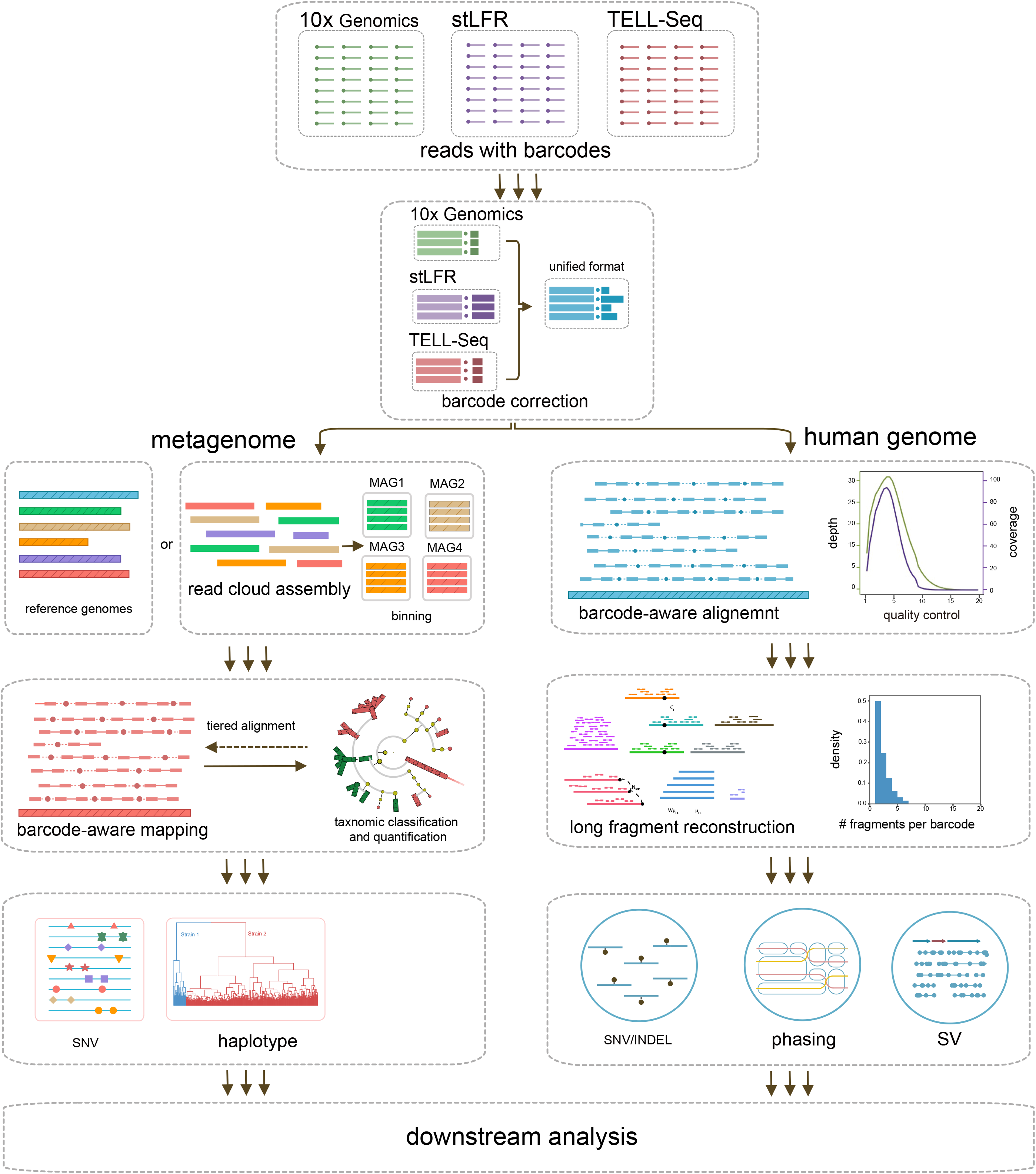
Overview of LRTK. The LRTK workflow includes a metagenomic section (left panel) and human genomic section (right panel). The metagenomic section implements barcode correction, read cloud assembly, barcode-aware alignment, long DNA fragment reconstruction, taxonomic classification and quantification, as well as SNV detection and phasing. The human genomic section implements barcode correction, barcode-aware alignment, long DNA fragment reconstruction, and detection and phasing of SNVs, INDELs and SVs.

In the human genome section, LRTK directly maps reads to the human genome using the modified EMA[12] approach and marks the duplicates for each barcode. LRTK reconstructs long DNA fragments through greedy extension based on the read alignment coordinates if the co-barcoded reads are within a specified distance[13]. After mapping, LRTK provides user a choice of several well-known tools for variant detection, including FreeBayes[14], SAMtools[15], GATK[16] for calling small variants; and Aquila[17], LinkedSV[18] and VALOR2[19] for calling large variants. For variant phasing, LRTK utilizes HapCUT2[20] and WhatsHap[21] to call phasing blocks **(Table S2)**. Further details are available in the **Method** section.

### LRTK support multiple linked-read sequencing technologies

LRTK can handle linked-read data from various sequencing technologies, including but not limited to 10x Genomics, stLFR, or TELL-Seq. Initially, LRTK converts the raw linked-read from 10x Genomics, stLFR and TELL-Seq into a unified FASTQ format, which contains a new field “BX:Z:” to store 16 bp (10x Genomics linked-read), 18 bp (TELL-Seq) and 30 bp (stLFR) barcode sequences **(Figure S1)**.

Because barcode sequences are typically located at the beginning or end of linked-reads, LRTK includes functions to correct potential sequencing errors. For 10x Genomics linked-read, after correction, there are approximately 94.8% of barcode sequences on whitelist for NA12878 and 94.1% for ATCC-MSA-1003, respectively. The performance is comparable to the results obtained from Long Ranger, which retains 94.4% of barcode sequences for NA12878 and 93.6% for ATCC-MSA-1003. The retention rates were slightly lower for stLFR linked-read, with 85.4% for NA12878 and 90.7% for ATCC-MSA-1003. A known concern with stLFR linked-read sequencing is the loss of barcode specificity during analysis. This occurs when the data is typically converted into a 10x-compatible format to run Long Ranger or directly aligned using BWA[22] without considering the barcode information[3]. We made modifications to EMA[12] **(Methods)** to accommodate barcodes with varying lengths, including but not limited to stLFR. Our adapted approach proved to be more effective at preserving barcode specificity than the 10x-compatible format conversion approach.

### LRTK supports reconstruction of long DNA fragments

The quality of DNA library may significantly affect assembly performance and variant calling[23]. To evaluate the quality of linked-read sequencing libraries, we reconstructed the long DNA fragments for both human genome and metagenome sequencing data. We also calculated several key statistics to comprehensively compare the fragment properties across different linked-read sequencing technologies. The statistics include average coverage of short reads per fragment (C_R_), average physical coverage of the genome by long DNA fragments (C_F_), number of fragments per partition (N_F/P_), fragment length (μ_FL)_, and unweighted and length-weighted DNA fragment length (μ_FL_ and Wμ_FL_) **(Figure S2)**. On human genome sequencing data (B2, NA12878), LRTK detected approximately 7.36, 1.22 and 3.09 fragments per barcode and achieved average fragment lengths of 50.19, 62.28 and 80.04 kb for 10x Genomics linked-read, stLFR and TELL-Seq, respectively (**Figure 2B**). On the metagenome sequencing data (ATCC-MSA-1003), stLFR linked-read yielded the lowest N_F/P_ (N_F/P_ =1.54), while TELL-Seq linked-read yielded a slightly higher number (N_F/P_ =4.26). Both numbers were much lower than that obtained from 10x Genomics linked-read (N_F/P_ =16.61) (**Figure 2A**), indicating that stLFR and TELL-Seq have superior performance in terms of the barcode specificity.

**Figure 2.**
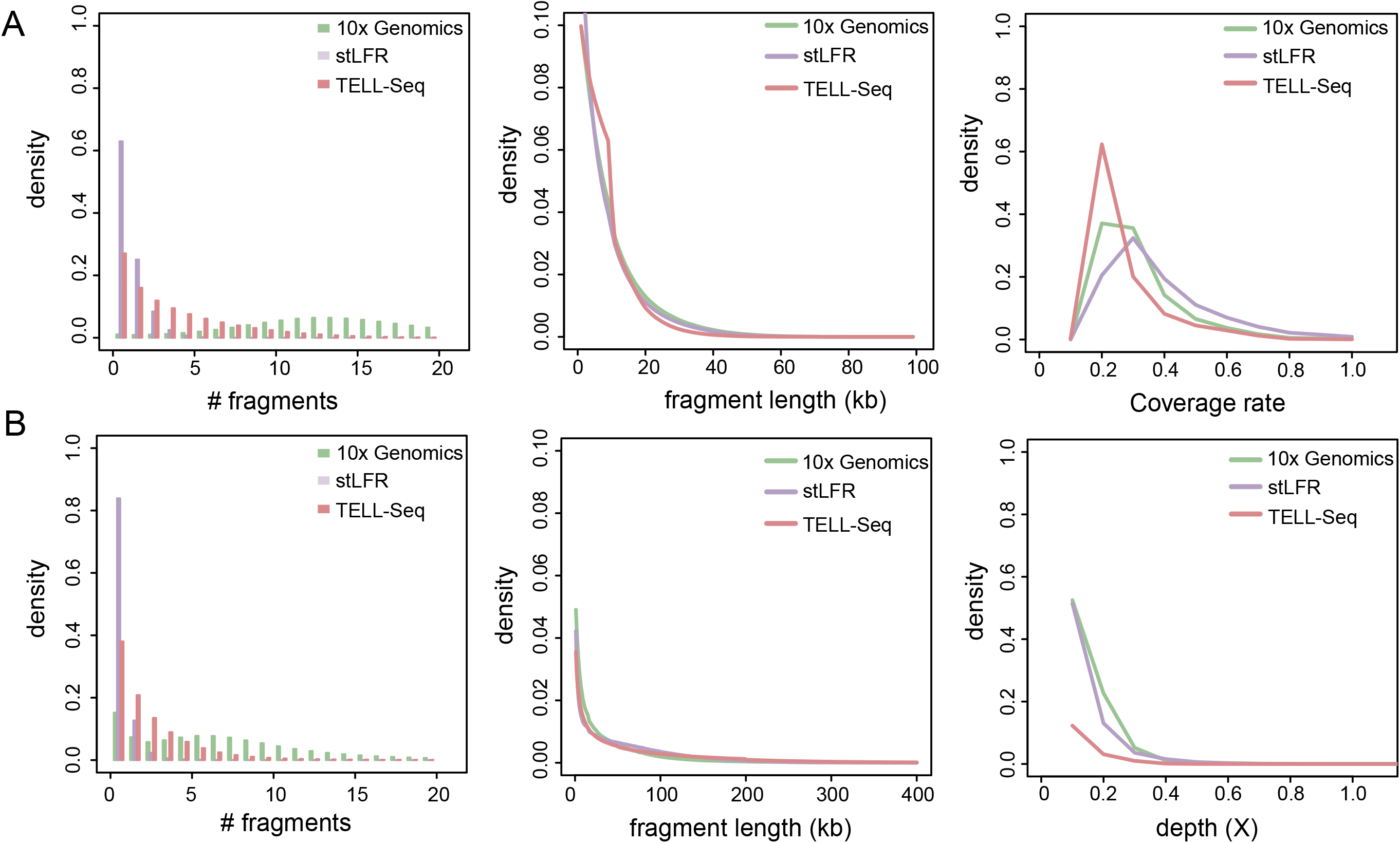
Quality metrics for different linked-read sequencing platforms. **(A)** Quality metrics under metagenome sequencing. The left panel displays the number of fragments per barcode for 10x Genomics, stLFR and INDEL in metagenomic sequencing data, respectively. The middle panel shows the DNA length of reconstructed fragments. The right panel shows the average read coverage per fragment. **(B)** Quality metrics under human genome sequencing. The left panel shows the number of fragments per barcode, while the middle panel shows the DNA length of reconstructed fragments. The right panel shows the average read coverage per fragment.

### LRTK supports metagenome variant detection and phasing using barcode-aware alignment

Previous studies have amply demonstrated the huge potential of linked-reads in metagenomics[5–9]. Here, we also evaluated the performance of linked-reads in detecting taxonomic abundance and metagenomic variants. Based on standardized microbial genome databases such as GTDB[24] and UHGG[11], we developed a reference genome based computational framework to identify species and SNVs from metagenomic sequencing data. Using the ATCC-MSA-1003 benchmarking data[23], we observed that LRTK had a comparable performance with *k-mer* based tools such as KMCP[25] and marker-gene based tools, MIDAS2[26] (**Figure 3A**). For high abundance species, LRTK could accurately identify SNVs from the alignment files and use barcode information to estimate genetic linkages and infer potential haplotypes. Comparison of metagenome SNV calling tools in the LRTK pipeline showed high consistency among FreeBayes[14], inStrain[27] and SAMtools[15] (**Figure 3B**). We applied LRTK on a real longitudinal linked-read metagenomic data set (D1)[6] to evaluate LRTK’s performance in practice. LRTK revealed the taxonomical composition and genomic variations for each sample. Using multiple related samples, LRTK also identified a genome-wide mirrored allele imbalance (MAI) for species *Alistipes finegoldii*, which may suggest intra-host strain evolution over time (**Figure 3C and D**).

**Figure 3.**
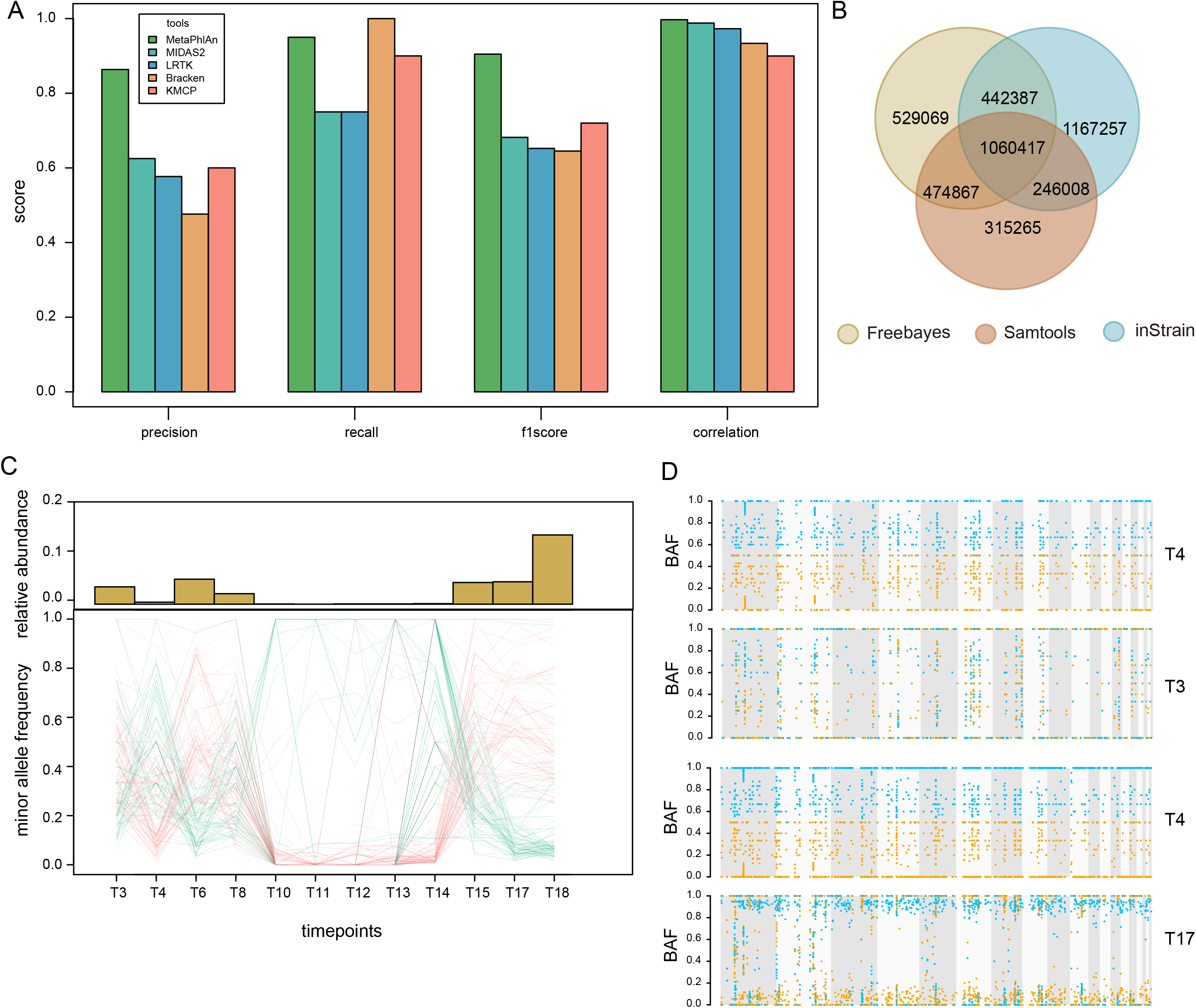
Comparison of linked-read based metagenomic quantification, SNV identification and phasing. **(A)** Evaluation of tools to quantify taxonomic abundance based on the linked-read metagenomic sequencing data. **(B)** Comparison of tools to detect SNVs. **(C)** Dynamic changes of taxonomic abundance and allele frequency. **(D)** SNV based strain phasing in pairwise samples.

### LRTK promotes metagenome assembly using barcode specificity

*De novo* assembly approaches have reconstructed numerous novel microorganisms from metagenomic sequencing data. Linked-read sequencing enhances metagenome assembly performance, especially for low abundance species[28]. Using benchmarking metagenomic sequencing data (ATCC-MSA-1003), we compared the performance of three linked-read based assemblers: Athena[28], Supernova[29] and Pangaea[30]. Among them, Pangaea achieved the best reconstructed genome fractions, NGA50, and NA50, for both stLFR and TELL-Seq sequencing data (**Figure 4A**). With Pangaea, LRTK achieves NA50 values of 1.8 Mb and 1.2 Mb for stLFR and TELL-Seq sequencing data, respectively. On 10x Genomics sequencing data, Athena exhibited superior assembly performance, with a NGA50 of 245 Kb. We applied LRTK on a dataset of human gut microbiomes (D2)[30] for illustrative purposes. Two of the assembled contigs were circularized genomes that showed near perfect collinearity with the closest reference genomes (**Figure 4B**). LRTK could also automatically clusters the contigs into bins after assembly. LRTK recovered 24 near-complete, 7 high quality and 52 medium quality bins, respectively (**Figure 4C-E**). The superior assembly performance we have observed affirms the effectiveness of linked-read sequencing technologies in metagenome assembly. Interestingly, the assembled metagenomes, due to their high quality, have the potential to serve as reference genomes. This characteristic makes LRTK valuable to longitudinal studies relying on linked-read metagenomic sequencing data.

**Figure 4.**
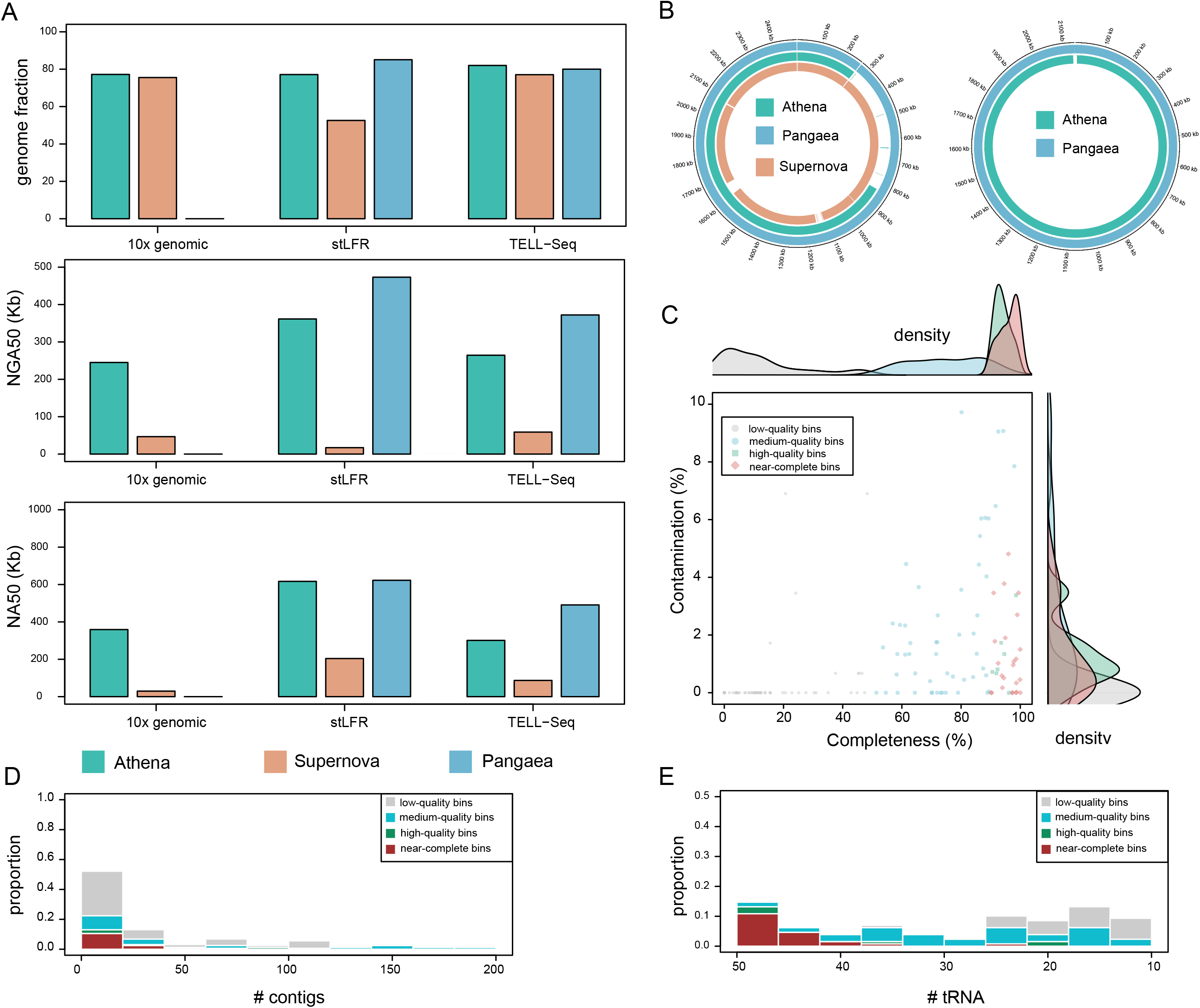
Evaluation of linked-read based metagenomic assembly. **(A)** Evaluation of assembly performance for Athena, Supernova and Pangaea using 10x Genomics, stLFR and TELL-Seq sequencing. **(B)** Illustration of the two assembled circular contigs. **(C)** The distribution of completeness and contamination for reconstructed bins. **(D)** The number of contigs in each contig group. **(E)** The number of detected tRNAs in each contig group.

### LRTK improves human genome variant phasing using long-range information

Previous studies have shown that the high base accuracy and long DNA fragments arising from linked-read sequencing benefits phasing of human genomic variants[31]. Here, we benchmarked the computational tools for variant detection and phasing tasks, and constructed a best practice to apply linked-reads on human genome studies. To ensure fair comparisons, LRTK used the modified version of the barcode-aware alignment tool, EMA[12], to map linked reads from different sequencing technologies to the human genome (GRCH38). We first benchmarked the commonly used tools FreeBayes[14], GATK[16] and SAMtools[15] to detect SNV and small insertions and deletions (INDEL). Among them, GATK achieved the best F1 score in detecting SNVs, followed by SAMtools and FreeBayes (**Figure 5A**). For INDEL calling, all three tools showed similar precision levels of around 0.5 while GATK had a better recall rate (**Figure 5B**). We compared the linked-read phasing tools, Whatshap[21] and HapCUT2[20], in combination with the aforementioned variant detection tools. We observed that HapCUT2 (**Figure 5C and D**) achieves longer phasing blocks and higher phased heterozygous SNV rate compared to WhatsHap[21]. Since most structural variation (SV) detection tools were developed to be used with 10x Genomics sequencing data and may therefore have uncertain performance loss when used with other sequencing technologies, we only benchmarked the performance of SV calling tools using 10x Genomics sequencing data. As illustrated in **Figure 5F and G**, Aquila had higher recall rates for 50 bp – 1 Kb deletions and insertions, while LinkedSV[18] performed better in detecting deletions longer than 1 Kb. We also applied LRTK on a family trio’s genomic sequencing dataset (D3: NA24143, NA24149 and NA24385) to evaluate its actual performance. LRTK demonstrated excellent phasing performance and improved identity-by-descent (IBD) segments detection for pairwise samples (**Figure 5E**).

**Figure 5.**
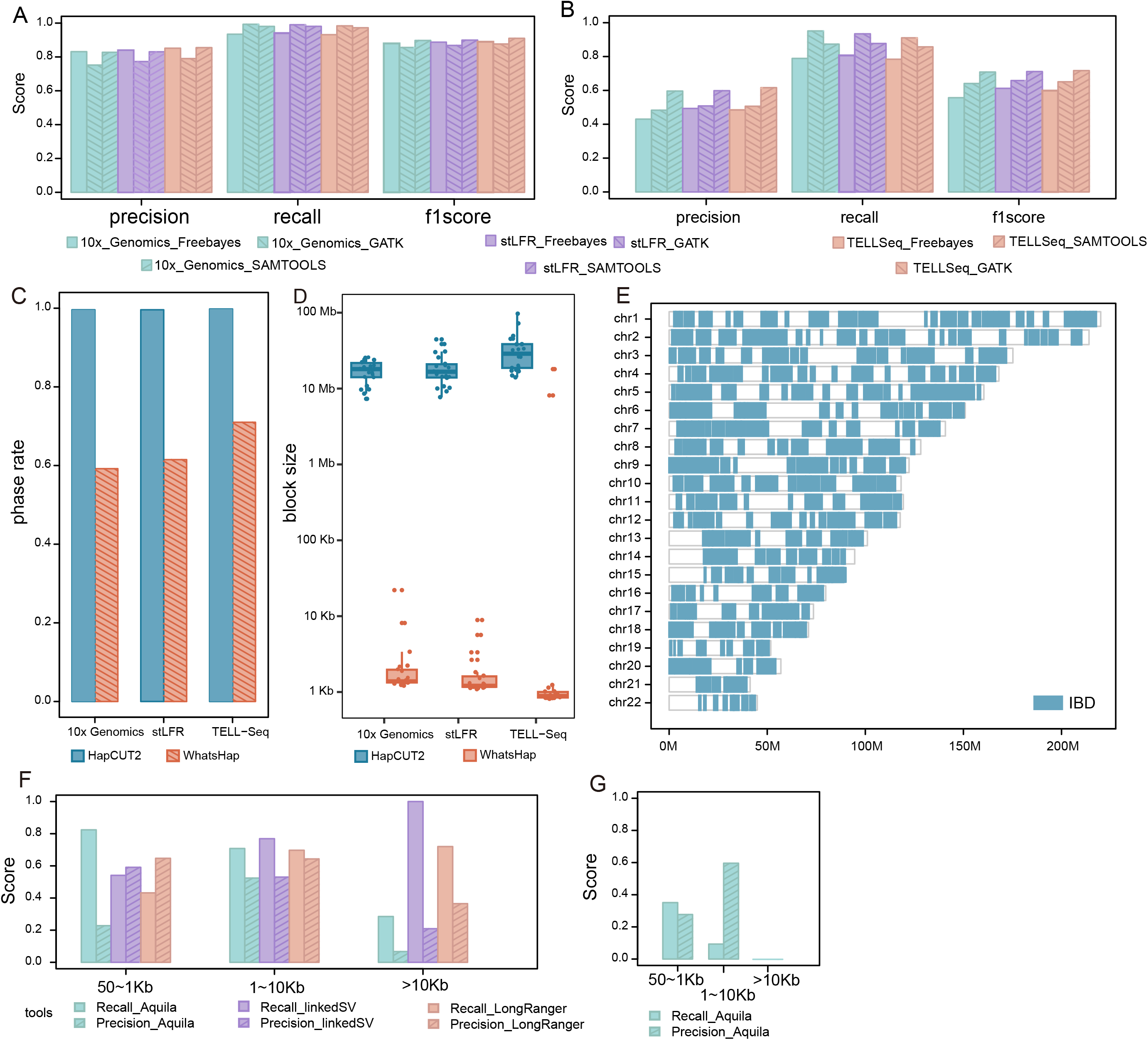
Evaluation of linked-read based detection of variation in the human genome. **(A, B)** Performance metrics on the detection of SNVs and INDELs using FreeBayes, GATK and SAMtools for 10x Genomics, stLFR and TELL-Seq. **(C, D)** The performance on phasing of small variants using HapCUT2 and WhatsHap for 10x Genomics, stLFR and TELL-Seq. **(E)** Illustration of the improved performance on IBD detection. **(F, G)** The performance on detection of large deletions **(F)** and insertions **(G)** using Aquila, LinkedSV and LongRanger.

### LRTK provides flexible commands to process sequencing data

A primary advantage of LRTK is its flexible, user-defined settings for different tasks. Users have a choice to run each LRTK module independently and generate separate results for each module. For instance, the MKFQ function could be independently used to simulate sequencing reads from the stLFR platform. For some functions, LRTK provides multiple tools for users to choose from. For metagenome SNVs detection, LRTK offers the user a choice between SAMtools[15], FreeBayes[14], and inStrain[27]. To accommodate user-specific requirements on LRTK command, LRTK also allows users to set different parameters and save the results for further comparison.

### LRTK provides automated analysis and user-friendly reports

LRTK provides an automated analysis pipeline, starting from raw sequencing reads to performing diverse data analysis and generating publication-ready visualizations. Specifically, LRTK produces different types of data features, calculates their corresponding statistical indicators, and presents them together in a HTML report. Taking the aforementioned longitudinal linked-read sequencing dataset D1 as an example, LRTK generates a systematic summary for the input data and analysis results of each step. Firstly, LRTK produces summary statistics for the FASTQ files obtained from the read quality control (QC) tools **(Figure S3A-B)**. After aligning reads to reference genomes, LRTK calculates the key parameters for library preparation and reconstructed long fragments **(Figure S3C-D)**. For detected species and variants, LRTK generates basic statistics and presents them using concise distribution plots **(Figure S3E-F).** For downstream analysis, LRTK conducts principal components analysis (PCA) analysis on the relative abundance profiles and clustering analysis on allele frequency to make comparisons across multiple samples **(Figure S3G-H)**. Similar reports could be produced for users employing LRTK to study the human genome **(Figure S4)**.

## Discussion

Seeing that multiple linked-read sequencing technologies are extensively utilized in scientific studies, a platform agnostic linked-read processing tool would be an intuitive solution to ensure reproducibility and robustness. Unfortunately, a cross-platform software solution is currently unavailable to the research community. Accordingly, we introduce LRTK, a unified and versatile computational framework to efficiently process sequencing data from 10x Genomics, stLFR and TELL-Seq technologies. LRTK includes separately invokable commands to perform linked-read simulation, barcode correction, barcode-aware alignment, read cloud based assembly reconstruction of long fragments and other barcode-assisted genomic variant calling and phasing. LRTK also provides automated and complete analysis, from raw data QC through advanced downstream analysis to generation of publication-ready visualization. LRTK is also open source, allows easy integration with other scientific pipelines.

One of the core advantages of LRTK is that it provides solutions for both metagenomic analysis and human genome analysis. Linked-read sequencing technologies offer highly accurate reads and inferable long-range information, which confers distinct advantages in genome research, such as superior phasing ability[31]. Phasing is valuable in exploring intra-host evolution of microbiota where strains under the same species may have some variants. LRTK follows a reference genome based approach to detect species’ abundance and to identify metagenome variants. It also infers the mirrored allele imbalance from multiple related samples. In addition, we establish a best practice to perform variation calling, phasing, and detection of IBD segments for human genome.

Finally, it is worth mentioning that LRTK has the potential to be extended to handle other types of linked-read sequencing technologies. For example, in 2017, Illumina introduced the bead-based barcode partitioning in a single tube to phase human genomes. It further proposed the complete long-read technology for complex genomes in 2022[32]. Additionally, Meier J I, *et al.* developed Haplotype tagging to investigate the butterfly species[33]. Redin D, *et al.* recently introduced a novel library preparation method for high throughput barcoding of short reads[34]. The new single cell metagenomic sequencing technologies employs highly accurate barcoded reads and provides inferable long-range information, which could potentially be used in combination with current linked-read technology in future studies[35]. We are actively developing LRTK to incorporate these technologies.

## Methods

### Data collection

We curated one benchmarking dataset and two real datasets to demonstrate the performance of LRTK in processing metagenomic sequencing data **(Table S3)**. The benchmarking datasets (B1: ATCC-MSA-1003) were obtained from the NCBI with the following accession numbers: SRR12283286 for 10x Genomics, and PRJNA875547 for stLFR and TELL-Seq sequencing technologies. The first real metagenomic dataset (D1), consisting of longitudinal 10x Genomics linked-read sequencing data, was downloaded from the NCBI under accession number SRP323279. We used only a subset of samples containing the barcode information in the read ID **(Table S3)**. Another real metagenomic dataset (D2) containing deep stLFR sequencing data, was downloaded from the China National GeneBank (CNGB) under project CNP0003432.

For the human genome section, we obtained the linked-read of NA12878 for 10x Genomics, stLFR and TELL-Seq and used them as the benchmarking dataset B2. We also downloaded the 10x Genomics linked-read sequencing data for a family trio (NA24143, NA24149 and NA24385) and used them as the real data set D3 **(Table S3)**.

### Data preprocessing

We utilized LRTK to convert the raw linked-read from 10x Genomics, stLFR and TELL-Seq into a unified FASTQ format **(Figure S1)** and correct potential sequencing errors in barcodes. For the 10x Genomics and stLFR linked-read, the barcodes are aligned to their respective barcode whitelists using the BWA aln command. Aligned barcodes containing fewer than 2 mismatches are then corrected as the corresponding barcodes in the whitelist. Due to the lack of a barcode whitelist for TELL-Seq, LRTK adopts the approach described by Chen *et al.*[3] to correct barcode errors. LRTK counts the supporting read for each barcode, compares barcodes with one supporting read to those with multiple supporting reads, and corrects potential sequencing errors for barcodes with one mismatch to those with multiple supporting reads. The linked-read in the unified FASTQ file are then subjected to fastp[36] to remove adapter sequences and low-quality reads. For metagenomic sequencing data, the sequencing reads are additionally aligned to the human genome and only unmapped microbial reads are retained. LRTK also provides support for simulating linked-read from 10x Genomics and stLFR platforms through modification of LRTK-SIM[23].

### Metagenome assembly and binning

We evaluated the performance of three different metagenome assemblers: Athena[28], Pangaea[30] and Supernova[29], and observed superior performance of Pangaea on stLFR and TELL-Seq sequencing data. Therefore, for LRTK, we chose Pangaea as the default assembler to assemble linked metagenomic sequencing data. After the initial assembly, we extracted the contigs that were at least 1Mb in length and checked their circularization. The uncircularized contigs were grouped using MetaBAT2[37].

### Barcode-aware read alignment

For human genome sequencing data, we utilized EMA, a barcode-aware alignment approach, to map high-quality reads to the human reference genome[38]. We modified EMA to be compatible with the barcodes from stLFR (30 bp) and TELL-Seq (18 bp). LRTK marks PCR duplicates for each barcode using the “BARCODE_TAG” parameter in Picard (https://broadinstitute.github.io/picard/). The alignment files are then sorted according to the genomic coordinates for further analysis.

For metagenomic sequencing data, we developed a tiered alignment approach to align the linked read to microbial genome using modified EMA. The default reference genomes for the human gut metagenome were downloaded from UHGG[11]. After the first alignment, we partitioned the reference genomes into 1-kb windows and calculated the number of mapped reads for each window. The mapped reads are categorized into two types: unique mapped reads (U) and reads with multiple alignments (M). Thus, we can use the following formulas to determine the total read count of the window (RC(W)), unique mapped read count (RC(U)) and multiply mapped read count (RC(M)), where I is the window size.

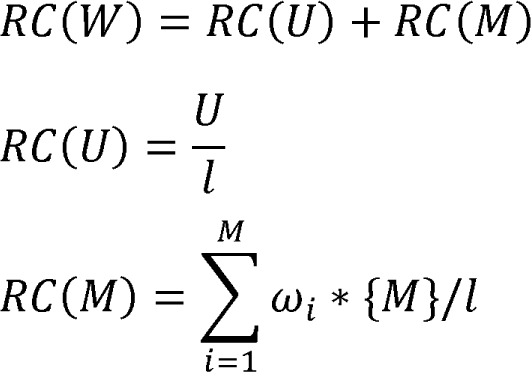

In the case of read with multiple alignments, a coefficient w is introduced. When a read in M has alignments with N different windows, w is calculated using the following formula:

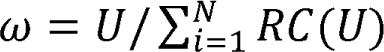

To ensure accurate estimation of genome abundance, we calculate it as the mean of the n% most closely covered windows. After initial classifying of microbial genomes, we extract their respective genome sequences and realign the reads to the new subset reference genome sequence Finally, we recalculate the relative abundance by using the most closely covered windows again.

### Reconstruction of long DNA fragments

We developed a computational strategy to reconstruct long DNA fragments from linked-read sequencing data. Initially LRTK extracts paired reads that share the same barcode in the alignment file. The uniquely aligned paired-end reads are collected to evaluate the distribution of insert sizes (with a mean of 1PE and a standard deviation of aPE). Alignments are removed if the distance between two reads in a pair (R1 and R2) exceeds a certain threshold, that is,

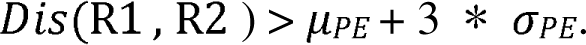

The remaining paired reads are used as seeds and extended to both directions to connect with other seeds sharing the same barcode until no more eligible seeds can be found within a specified distance. All of these co-barcoded reads are considered to be derived from the same long DNA fragment and are collected as a read cloud.

### Identify and phase genomic variants

LRTK supports multiple popular variant detection tools and adheres to best practices for linked-read sequencing. For human genomic sequencing data, LRTK provides FreeBayes[14], SAMtools[15] and GATK[16] to call SNVs and INDELs. The available phasing tools include HapCUT2[20] and WhatsHap[21]. Having obtained the phased SNVs, we then use hap-ibd[39] to detect pairwise identity-by-descent segments across multiple samples. By default, large structural variations (SVs) are identified and phased using Aquila[17]. However, Users have the flexibility to select other tools such as LinkedSV[18] and VALOR2[19] for SV analysis.

For metagenome sequencing data, LRTK offers three metagenome SNV callers: FreeBayes[14], inStrain[27] and SAMtools[15]. The SNV phasing is performed on high abundance species using WhatsHap[21] with the inferred ploidy. LRTK also selects SNVs located on certain high abundance species and compares the SNV profiles across multiple samples. Based on the allele frequency of SNVs, we used an unsupervised clustering method to detect potential minor allele imbalance (MAI) events for pairwise samples. In brief, one sample is chosen as the reference for all the contigs. The minor allele frequency of heterozygous SNVs present in others samples are then compared to the reference sample. We used the k-means clustering method to separate the SNVs into distinct groups using the minor allele frequency matrix. The number of clusters were determined by using the Calinski-Harabasz index. The number of clusters may be used to determine the number of strains under the same species.

### Downstream analysis and HTML-based visualization

The human genome analysis report begins with generating the FASTQ quality control statistics during the preprocessing step. For the barcode aware alignment step, it presents the distribution of several key statistics about DNA library, including the number of fragments per barcode, fragment length and read coverage per fragment. It also includes information about the sequencing coverage and insert size. The variant calling step summarizes the number of SNVs, small INDELs and large SVs and illustrates their distributions.

Similarly, the metagenome analysis report firstly illustrates the QC results for preprocessing, alignment and variant calling steps. Additionally, it includes an optional report about the automatic analysis of multiple related samples. The report shows the distribution of high abundant species across multiple samples, and uses a principal component analysis (PCA) plot to visualize the divergence across them. The report also depicts the distribution of SNVs of these high abundant species, and the minor allele frequency distribution in pairwise samples.

### Code Availability and Requirements

Source code is available at https://github.com/ericcombiolab/LRTK. LRTK can be downloaded as a packaged conda environment (https://anaconda.org/bioconda/lrtk).

Project name: Linked Read ToolKits project

Project home page: https://github.com/ericcombiolab/LRTK Operating system(s): Linux and macOS

Programming language: C and python

Other requirements: Conda, Python 3.6 or higher License: MIT

RRID: SCR_023945

### Data Availability

We curated two benchmarking datasets and three real datasets to evaluate the performance of LRTK in processing linked-read sequencing data.

**B1:** The benchmarking dataset B1 contains the 10x Genomics, stLFR and TELL-Seq sequencing data for ATCC MSA-1003 mock community. These datasets were obtained from the NCBI with the following accession numbers: SRR12283286 for 10x Genomics, and PRJNA875547 for stLFR and TELL-Seq sequencing technologies.

**B2:** The benchmarking dataset B2 contains the 10x Genomics, stLFR and TELL-Seq sequencing data for human sample NA12878. The linked-read of for 10x Genomics and stLFR were downloaded from GIAB while the raw TELL-Seq data was downloaded with the accession number SRR10584152 (PRJNA591637).

**D1:** The first real metagenomic dataset (D1), consisting of longitudinal 10x Genomics linked-read sequencing data, was downloaded from the NCBI under accession number SRP323279. We used only a subset of samples containing the barcode information in the read ID.

**D2:** The second real metagenomic dataset (D2), containing deep stLFR sequencing data, was downloaded from the China National GeneBank (CNGB) under project CNP0003432.

**D3:** We also downloaded the linked-read sequencing data for a family trio (NA24143, NA24149 and NA24385) and used them as the real data set D3 (Table S3). The 10x Genomics and stLFR sequencing data was downloaded from GIAB. For TELL-Seq sequencing data, we only obtained the raw TELL-Seq data from NA24385 from SRA database (PRJNA591637: including SRR10689414, SRR10689415, SRR10689416, SRR10689417).

## CRediT authorship contribution statement

**Chao Yang:** Writing - original draft, and preparing figures and tables, preparing source codes; **Zhenmiao Zhang:** preparing source codes; **Xuefeng Xie:** Consolidating resources; **Yufen Huang:** Consolidating resources; **Herui Liao:** preparing source codes; **Werner P Veldsman:** Revising - original draft, and visualizations; **Jin Xiao:** Revising - original draft, and visualizations; **Kejing Yin:** Supervision; **Xiaodong Fang:** Supervision; **Lu Zhang:** Project administer, Writing – review & editing, Supervision, and funding acquisition.

## Acknowledgments

This research was partially supported by the open project of BGI-Shenzhen, Shenzhen 518000, China (BGIRSZ20220012), the Hong Kong Research Grant Council Early Career Scheme (HKBU 22201419), HKBU Start-up Grant Tier 2 (RC-SGT2/19-20/SCI/007), HKBU IRCMS (No. IRCMS/19-20/D02), the Guangdong Basic and Applied Basic Research Foundation (No. 2021A1515012226), the Science Technology and Innovation Committee of Shenzhen Municipality, China (SGDX20190919142801722) and Shenzhen Science and Technology Innovation Commission (SZSTI) - Shenzhen Virtual University Park (SZVUP) Special Fund Project (No. 2021Szvup135).

## Additional Files

Figure S1. LRTK text file specification.

Figure S2. Ideogrammatic definitions of CR, CF, NF/P, μFL, and WμFL metrics.

Figure S3. Demo reports for metagenomics sequencing.

Figure S4. Demo reports for human genome sequencing.

Table S1. Comparison of LRTK with other published linked-read pipelines.

Table S2. Bioinformatics tools included in LRTK.

Table S3. Linked-read sequencing data sources

## Abbreviations

stLFR: single-tube long fragment read
TELL-Seq: transposase enzyme linked long-read sequencing
SNV: single nucleotide variant
INDEL: small insertion and deletion
SV: structural variation

## Figures and Tables

**Supplementary Figure 1**: LRTK text file specification. (A) An example of the unified FASTQ format. Each read contains a barcode field “BX:Z:barcode” after the read names. The lengths of barcodes are 16 bp, 30 bp and 18 bp for 10x Genomics linked-read, stLFR and TELL-Seq, respectively. (B) An example of the BAM file generated by LRTK. The barcode information is stored in the “BX:Z:barcode” field.

**Supplementary Figure 2.** Ideogrammatic definitions of C_R_, _CF_, N_F/P, μFL, and WμFL met__r__ics. (A) CR: Average depth of short reads per fragment. (B) CF_: Average physical depth of the genome by long DNA fragmen_ts_. (C) N_F/P_: Numbe_r_ of fragments per barcode. (D) Length-weighted average (μ_FL_) and unweighted average (Wμ_FL_) of DNA fragment lengths.

**Supplementary Figure 3.** Demo reports for metagenomics sequencing. (A) Per base sequencing quality scores along the reads. (B) Per Base GC content along the reads. (C) The number of fragments per barcode. (D) Fragment length distribution. (E). The abundance of the top 10 most abundant species. (F) The number of SNVs per species. (G) A principal component analysis using sample data from different groups.

**Supplementary Figure 4:** Demo reports for human genome sequencing. (A) Per base sequencing quality scores along the reads. (B) Per Base GC content along the reads. (C) The number of fragments per barcode. (D) Fragment length distribution. (E) Sequencing depth frequency. (F) The distribution of sequencing depth along a whole genome. (G) The distribution of inferred insert sizes. (H) The distribution of detected deletions. (I) The distribution of detected insertions.

**Table S1:** Comparison of LRTK with other published linked-read pipelines.

|  | Items | Long Ranger | LRTK |
| --- | --- | --- | --- |
| **Input** | 10x Genomics | Y | Y |
|  | stLFR | N | Y |
|  | TELL-Seq | N | Y |
| **Modules** | Linked-read preprocessing | Y | Y |
|  | metagenome quantification | N | Y |
|  | metagenome SNV detection | N | Y |
|  | metagenome phasing | N | Y |
|  | metagenome assembly | N | Y |
|  | metagenome downstream analysis | N | Y |
|  | human genome alignment | Y | Y |
|  | human genome SNV detection | Y | Y |
|  | human genome SV detection | Y | Y |
|  | human genome phasing | Y | Y |
|  | human genome downstream analysis | N | Y |

**Table S2.**
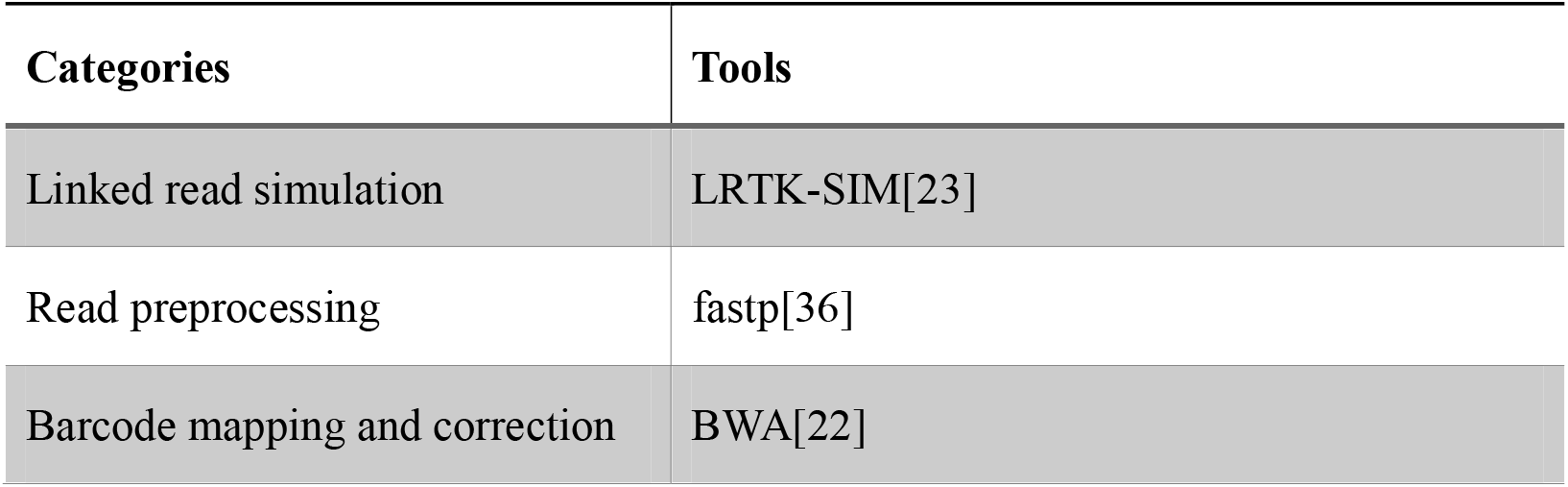
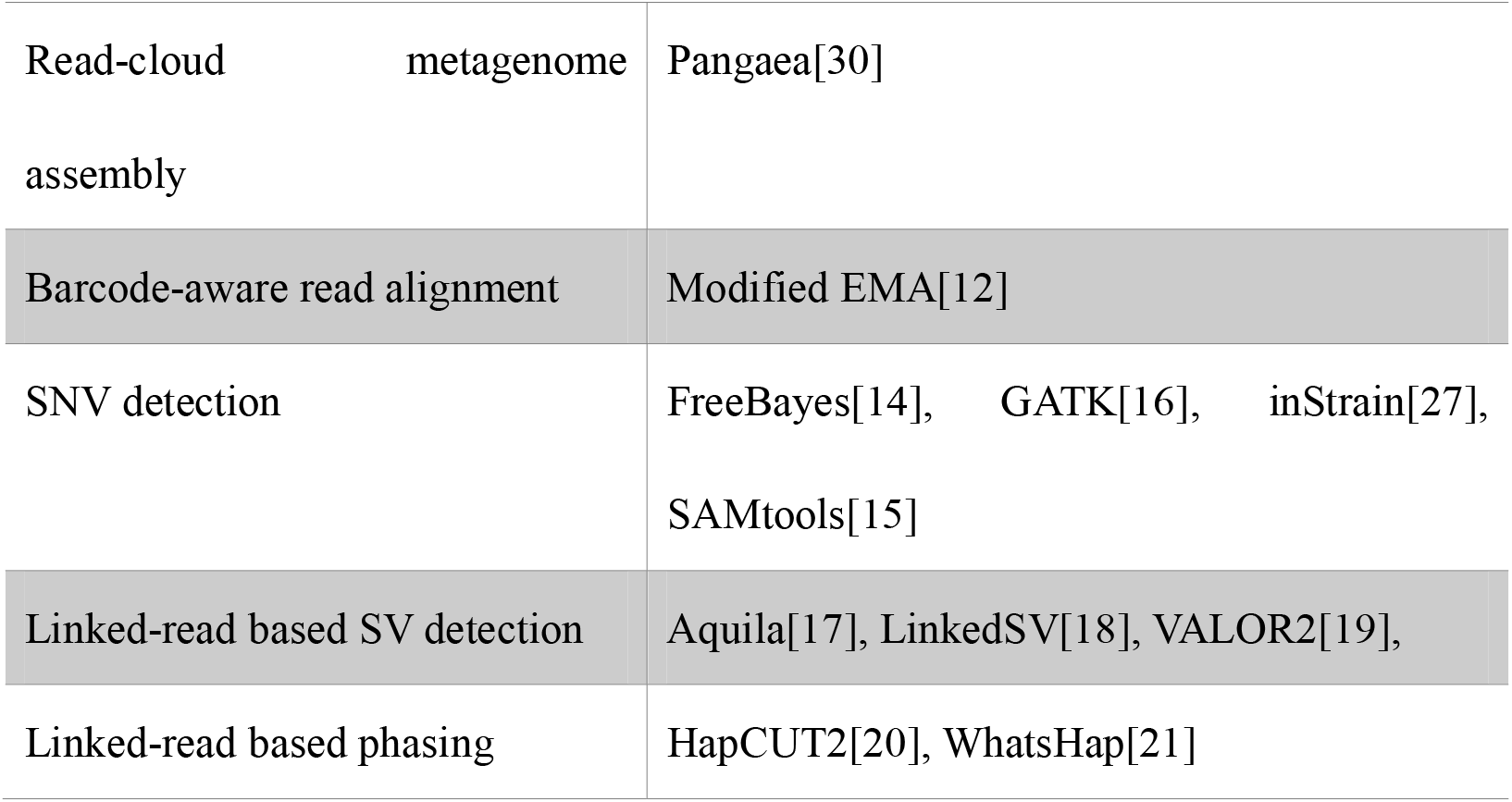
Bioinformatics tools included in LRTK.

**Table S3.**
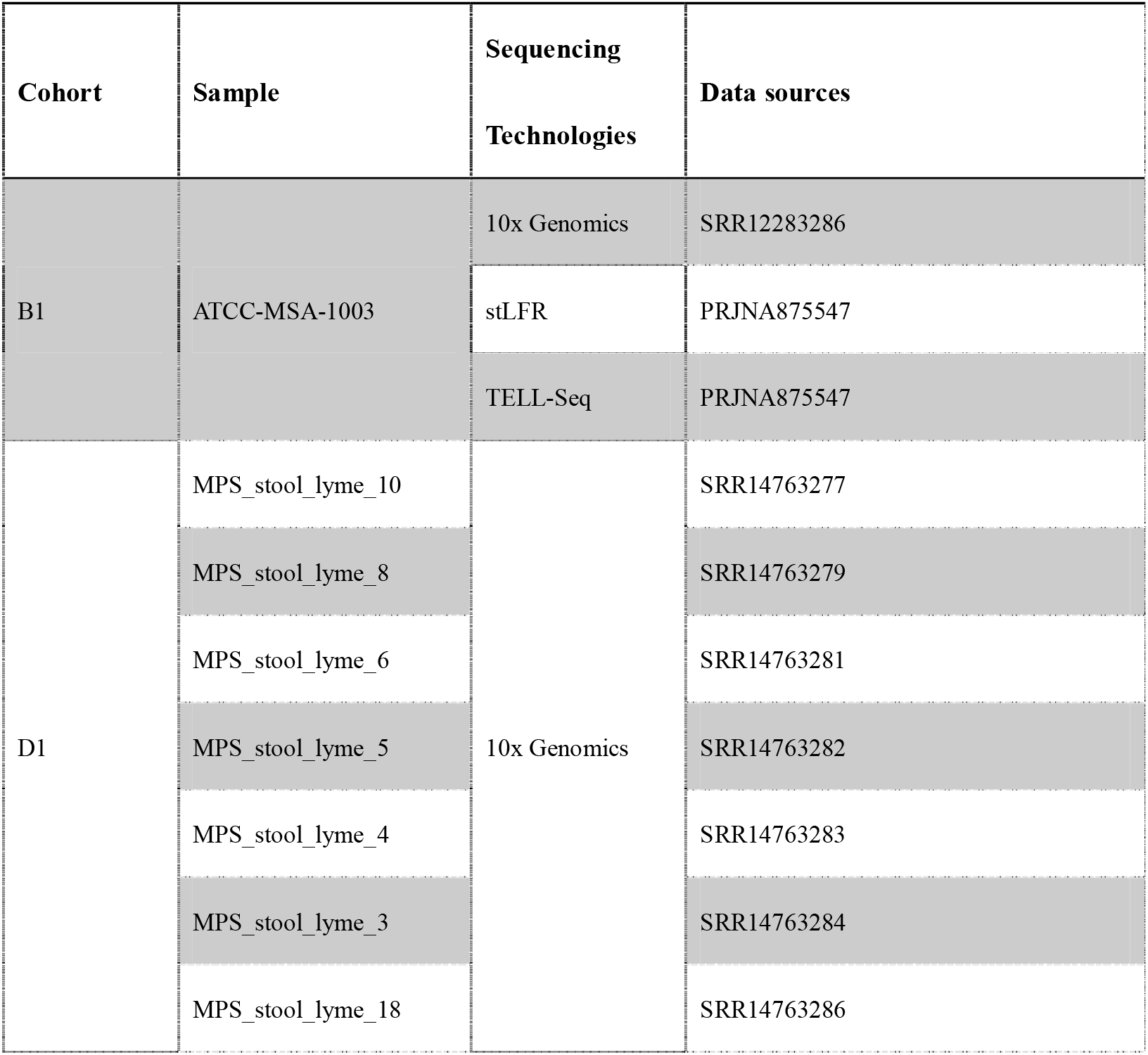
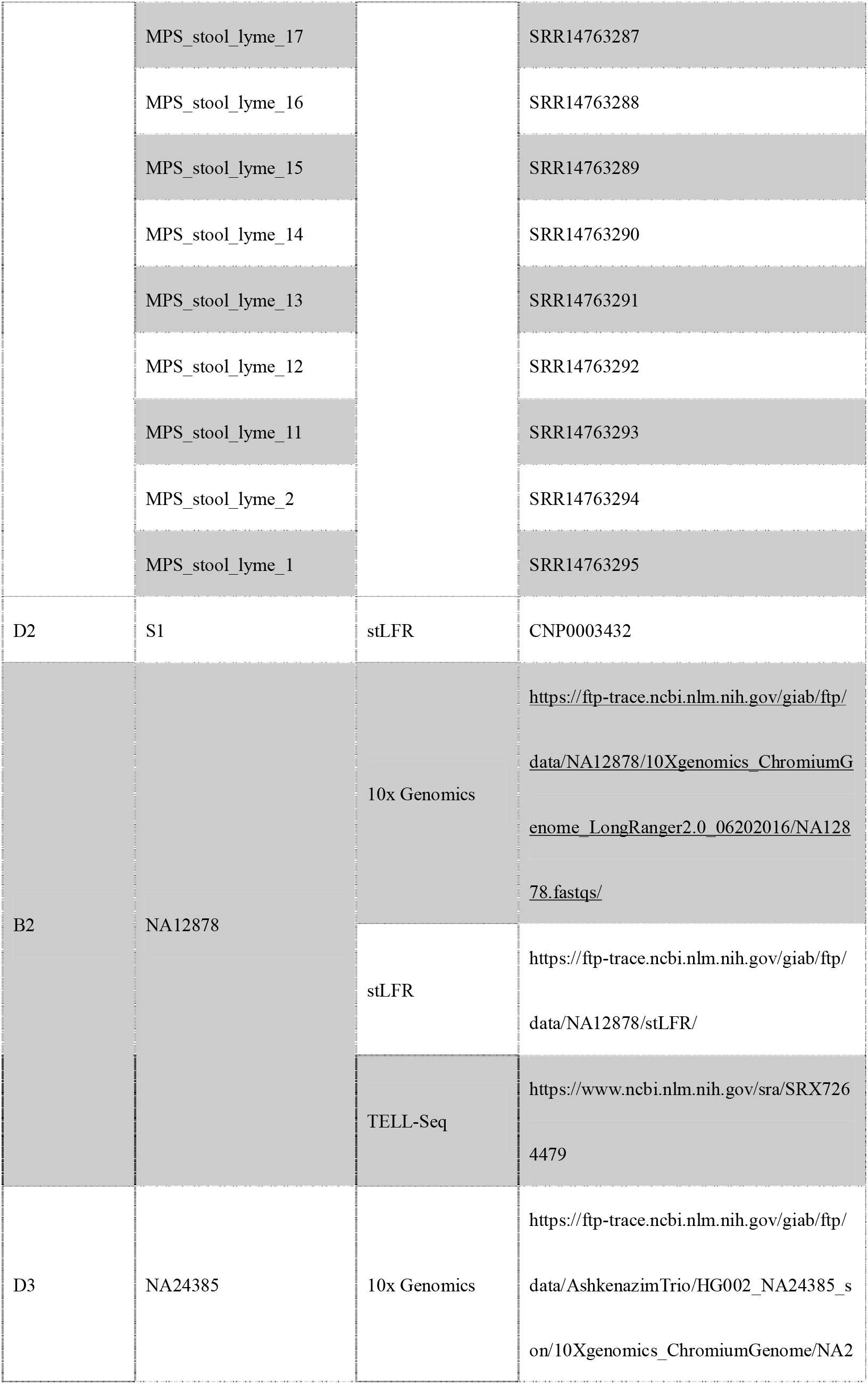
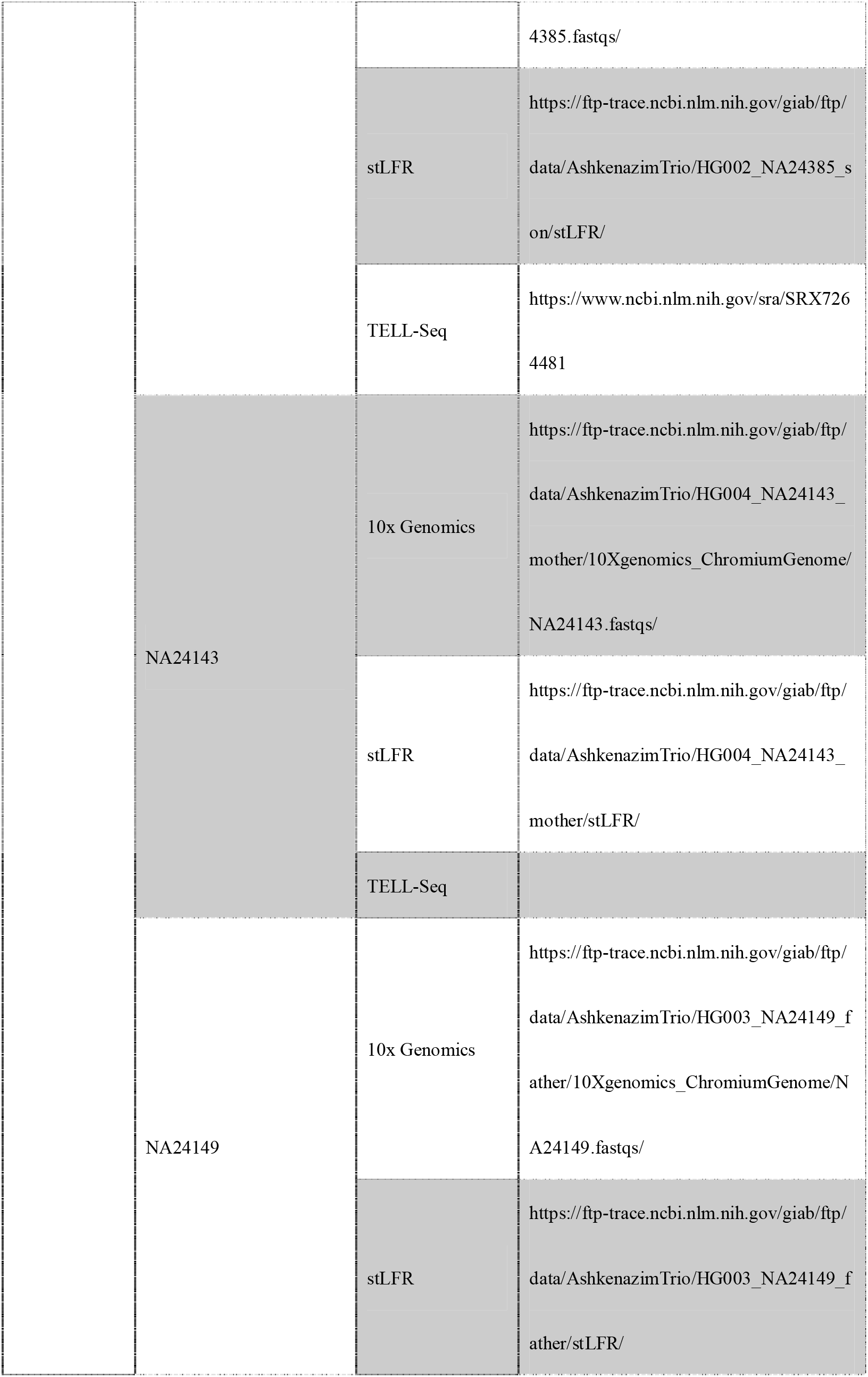
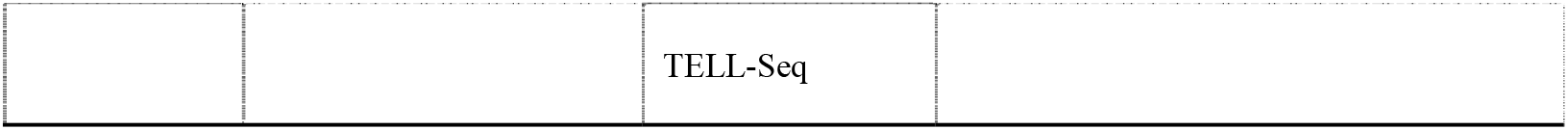
Linked-read sequencing data sources.

## References

1. Eisenstein M. Startups use short-read data to expand long-read sequencing market. Nat Biotechnol. Nat Biotechnol; 2015; doi: 10.1038/NBT0515-433.

2. Wang O, Chin R, Cheng X, Yan Wu MK, Mao Q, Tang J, et al.. Efficient and unique co-barcoding of second-generation sequencing reads from long DNA molecules enabling cost effective and accurate sequencing, haplotyping, and de novo assembly. Genome Res. Cold Spring Harbor Laboratory Press; 2019; doi: 10.1101/GR.245126.118.

3. Chen Z, Pham L, Wu TC, Mo G, Xia Y, Chan PL, et al.. Ultralow-input single-tube linked-read library method enables short-read second-generation sequencing systems to routinely generate highly accurate and economical long-range sequencing information. Genome Res. Cold Spring Harbor Laboratory Press; 2020; doi: 10.1101/gr.260380.119.

4. Spies N, Weng Z, Bishara A, McDaniel J, Catoe D, Zook JM, et al.. Genome-wide reconstruction of complex structural variants using read clouds. Nat Methods 2017 149. Nature Publishing Group; 2017; doi: 10.1038/nmeth.4366.

5. Siranosian BA, Brooks EF, Andermann T, Rezvani AR, Banaei N, Tang H, et al.. Rare transmission of commensal and pathogenic bacteria in the gut microbiome of hospitalized adults. Nat Commun 2022 131. Nature Publishing Group; 2022; doi: 10.1038/s41467-022-28048-7.

6. Roodgar M, Good BH, Garud NR, Martis S, Avula M, Zhou W, et al.. Longitudinal linked-read sequencing reveals ecological and evolutionary responses of a human gut microbiome during antibiotic treatment. Genome Res. Cold Spring Harbor Laboratory Press; 2021; doi: 10.1101/GR.265058.120/-/DC1.

7. Huang Y, Jiang P, Liang Z, Chen R, Yue Z, Xie X, et al.. Assembly and analytical validation of a metagenomic reference catalog of human gut microbiota based on co-barcoding sequencing. *Front Microbiol*. Frontiers; 2023; doi: 10.3389/FMICB.2023.1145315.

8. Davila Aleman FD. Microbiome and aging: A study of microbial evolution and community structure across model organisms. Abtract and Metadata. 2022;

9. Tracanna V, Ossowicki A, Petrus MLC, Overduin S, Terlouw BR, Lund G, et al.. Dissecting Disease-Suppressive Rhizosphere Microbiomes by Functional Amplicon Sequencing and 10× Metagenomics. mSystems. American Society for Microbiology; 2021; doi: 10.1128/MSYSTEMS.01116-20/SUPPL_FILE/MSYSTEMS.0116-20-S0001.PDF.

10. Zheng GXY, Lau BT, Schnall-Levin M, Jarosz M, Bell JM, Hindson CM, et al.. Haplotyping germline and cancer genomes with high-throughput linked-read sequencing. Nat Biotechnol 2016 343. Nature Publishing Group; 2016; doi: 10.1038/nbt.3432.

11. A A, S N, M B, F S, M B, ZJ S, et al.. A unified catalog of 204,938 reference genomes from the human gut microbiome. *Nat Biotechnol*. Nat Biotechnol; 2021; doi: 10.1038/S41587-020-0603-3.

12. Shajii A, Numanagić I, Whelan C, Berger B. Statistical Binning for Barcoded Reads Improves Downstream Analyses. Cell Syst. Cell Press; 2018; doi: 10.1016/J.CELS.2018.07.005/ATTACHMENT/8C829484-1F22-47F4-A513-B90845BD41D7/MMC1.PDF.

13. Zhang L, Zhou X, Weng Z, Sidow A. Assessment of human diploid genome assembly with 10x Linked-Reads data. *Gigascience*. Gigascience; 2019; doi:10.1093/GIGASCIENCE/GIZ141.

14. Garrison E, Marth G. Haplotype-based variant detection from short-read sequencing. 2012; doi: 10.48550/arxiv.1207.3907.

15. Li H, Handsaker B, Wysoker A, Fennell T, Ruan J, Homer N, et al.. The Sequence Alignment/Map format and SAMtools. Bioinformatics. Oxford Academic; 2009; doi: 10.1093/BIOINFORMATICS/BTP352.

16. McKenna A, Hanna M, Banks E, Sivachenko A, Cibulskis K, Kernytsky A, et al.. The Genome Analysis Toolkit: A MapReduce framework for analyzing next-generation DNA sequencing data. Genome Res. Cold Spring Harbor Laboratory Press; 2010; doi: 10.1101/GR.107524.110.

17. Zhou X, Zhang L, Weng Z, Dill DL, Sidow A. Aquila enables reference-assisted diploid personal genome assembly and comprehensive variant detection based on linked reads. Nat Commun 2021 121. Nature Publishing Group; 2021; doi: 10.1038/s41467-021-21395-x.

18. Fang L, Kao C, Gonzalez M V., Mafra FA, Pellegrino da Silva R, Li M, et al.. LinkedSV for detection of mosaic structural variants from linked-read exome and genome sequencing data. Nat Commun 2019 101. Nature Publishing Group; 2019; doi: 10.1038/s41467-019-13397-7.

19. KaraoClanoClu F, Ricketts C, Ebren E, Rasekh ME, Hajirasouliha I, Alkan C. VALOR2: characterization of large-scale structural variants using linked-reads. Genome Biol. BioMed Central Ltd.; 2020; doi:10.1186/S13059-020-01975-8/TABLES/2.

20. Edge P, Bafna V, Bansal V. HapCUT2: robust and accurate haplotype assembly for diverse sequencing technologies. Genome Res. Genome Res; 2017; doi: 10.1101/GR.213462.116.

21. Patterson MD, Marschall T, Pisanti N, Van Iersel L, Stougie L, Klau GW, et al.. WhatsHap: Weighted Haplotype Assembly for Future-Generation Sequencing Reads. https://home.liebertpub.com/cmb. Mary Ann Liebert, Inc. 140 Huguenot Street, 3rd Floor New Rochelle, NY 10801 USA ; 2015; doi: 10.1089/CMB.2014.0157.

22. Li H, Durbin R. Fast and accurate short read alignment with Burrows–Wheeler transform. Bioinformatics. Oxford Academic; 2009; doi: 10.1093/BIOINFORMATICS/BTP324.

23. Zhang L, Fang X, Liao H, Zhang Z, Zhou X, Han L, et al.. A comprehensive investigation of metagenome assembly by linked-read sequencing. Microbiome. BioMed Central Ltd; 2020; doi: 10.1186/S40168-020-00929-3/FIGURES/4.

24. DH P, M C, C R, AJ M, PA C, P H. GTDB: an ongoing census of bacterial and archaeal diversity through a phylogenetically consistent, rank normalized and complete genome-based taxonomy. *Nucleic Acids Res*. Nucleic Acids Res; 2021; doi: 10.1093/NAR/GKAB776.

25. Shen W, Xiang H, Huang T, Tang H, Peng M, Cai D, et al.. KMCP: accurate metagenomic profiling of both prokaryotic and viral populations by pseudo-mapping. Bioinformatics. Oxford Academic; 2023; doi: 10.1093/BIOINFORMATICS/BTAC845.

26. Zhao C, Dimitrov B, Goldman M, Nayfach S, Pollard KS. MIDAS2: Metagenomic Intra-species Diversity Analysis System. Bioinformatics. Oxford Academic; 2023; doi: 10.1093/BIOINFORMATICS/BTAC713.

27. Olm MR, Crits-Christoph A, Bouma-Gregson K, Firek BA, Morowitz MJ, Banfield JF. inStrain profiles population microdiversity from metagenomic data and sensitively detects shared microbial strains. Nat Biotechnol 2021 396. Nature Publishing Group; 2021; doi: 10.1038/s41587-020-00797-0.

28. A B, EL M, M K, AE P, Z W, A S, et al.. High-quality genome sequences of uncultured microbes by assembly of read clouds. *Nat Biotechnol*. Nat Biotechnol; 2018; doi: 10.1038/NBT.4266.

29. Visendi P. De Novo Assembly of Linked Reads Using Supernova 2.0. Methods Mol Biol. Humana Press Inc.; 2022; doi: 10.1007/978-1-0716-2067-0_12/COVER.

30. Zhang Z, Wang H, Yang C, Huang Y, Yue Z, Chen Y, et al.. Exploring high-quality microbial genomes by assembly of linked-reads with high barcode specificity using deep learning. bioRxiv. Cold Spring Harbor Laboratory; 2022; doi: 10.1101/2022.09.07.506963.

31. Chaisson MJP, Sanders AD, Zhao X, Malhotra A, Porubsky D, Rausch T, et al.. Multi-platform discovery of haplotype-resolved structural variation in human genomes. Nat Commun 2019 101. Nature Publishing Group; 2019; doi: 10.1038/s41467-018-08148-z.

32. Zhang F, Christiansen L, Thomas J, Pokholok D, Jackson R, Morrell N, et al.. Haplotype phasing of whole human genomes using bead-based barcode partitioning in a single tube. Nat Biotechnol 2017 359. Nature Publishing Group; 2017; doi: 10.1038/nbt.3897.

33. Meier JI, Salazar PA, Kučka M, Davies RW, Dréau A, Aldás I, et al.. Haplotype tagging reveals parallel formation of hybrid races in two butterfly species. Proc Natl Acad Sci U S A. National Academy of Sciences; 2021; doi: 10.1073/PNAS.2015005118/SUPPL_FILE/PNAS.2015005118.SD06.XLSX.

34. Redin D, Frick T, Aghelpasand H, Käller M, Borgström E, Olsen RA, et al.. High throughput barcoding method for genome-scale phasing. Sci Reports 2019 91. Nature Publishing Group; 2019; doi: 10.1038/s41598-019-54446-x.

35. Zheng W, Zhao S, Yin Y, Zhang H, Needham DM, Evans ED, et al.. High-throughput, single-microbe genomics with strain resolution, applied to a human gut microbiome. Science (80-). American Association for the Advancement of Science; 2022; doi: 10.1126/SCIENCE.ABM1483/SUPPL_FILE/SCIENCE.ABM1483_MOVIES_S1_TO_S10.ZIP.

36. Chen S, Zhou Y, Chen Y, Gu J. fastp: an ultra-fast all-in-one FASTQ preprocessor. Bioinformatics. Oxford Academic; 2018; doi: 10.1093/BIOINFORMATICS/BTY560.

37. DD K, F L, E K, A T, R E, H A, et al.. MetaBAT 2: an adaptive binning algorithm for robust and efficient genome reconstruction from metagenome assemblies. *PeerJ*. PeerJ; 2019; doi: 10.7717/PEERJ.7359.

38. Schneider VA, Graves-Lindsay T, Howe K, Bouk N, Chen HC, Kitts PA, et al.. Evaluation of GRCh38 and de novo haploid genome assemblies demonstrates the enduring quality of the reference assembly. Genome Res. Cold Spring Harbor Laboratory Press; 2017; doi: 10.1101/GR.213611.116.

39. Zhou Y, Browning SR, Browning BL. A Fast and Simple Method for Detecting Identity-by-Descent Segments in Large-Scale Data. Am J Hum Genet. Cell Press; 2020; doi: 10.1016/J.AJHG.2020.02.010.

