## Supplementary figures 1-4 and tables 1-3 for "LRTK: A platform agnostic toolkit for linked-read analysis of both human genomes and metagenomes": SupplementaryFigure1.pdf

**A.**

```
@ST-E00273:177:HMTTCCCXX:1:2110:11058:69133 BX:Z:AAACACCAGATGTTAG-1
AGTTGTTATGGGAAGATATTTCCTTTTTCAACATAGGCCTGAAAGCGCTCCAAATGTCCACTTCCAGATACTACAAAAGGAGTGATTCCAACCTGCTCTATGATAGGGAATGTTCAACTCTGTGTCCT
+
KKKKKKFKKFFKKKKKKKKKKKKKKKKFKKKKKKKKKKKKAKKKKKKKFFKKFKKKKKKKKKKKKKFKKKKKKKKKKKKKKKKFKKKKKKKKKKKKFFKKK,F7KF,AAKAFKKFAFKFKKAAFKAK,
@ST-E00273:177:HMTTCCCXX:1:2209:28747:51184 BX:Z:AAACACCAGATGTTAG-1
CTGTAACTTAAACAAATTTACAAGAAACAACAGCAACAACAACAAAAACATTAAAAAGCAGACAAAAGACATGAACAGGTACTTTTCAACAGAAGACATACATGTGGCCAACAATCACATGAGCAAAA
+
KKKKKKKKKKKKKKKKKKKKKKKKKKKKKKKKKKKKKKKKKKKKKKKKAKKKKKKKKKKKKKKKKKKKKKKKKKKKKFAKKKKKKKKKKKKKKKKKKKKKKKKKKKKKFFKFFFKKKKKKFAFKFKKKK
@ST-E00273:177:HMTTCCCXX:1:2210:24261:62611 BX:Z:AAACACCAGATGTTAG-1
AAGAATGTGTATTAGTTTACTTGCGTTTTTCCCTATATAAATCATTGGCAATGAGAAATGATAAACTCATTCTGCCCAGTATTATGTGGGTGATCAATAATTAGTATTAGAAAACAACTTTACAGTGC
+
KKKKKKKKKKKKKKKKKKKKKKKKKKKKKKKKKKKKKKKKKKKKKKKKKKKKKKKKKKKKKKKKKKKKKKKKKKKKKKKKKKKKKKKKKKKKKKKKKKKKKKKKKKKKKKKKKKKKKKKKKKKKKKKKA
@ST-E00273:177:HMTTCCCXX:1:2210:25022:61292 BX:Z:AAACACCAGATGTTAG-1
ATGGATGAATGAATGCATGAACATATGGATGGATGGATAAGTGAATTGATGACTGGATGGATGGTTGAATGGATGGATAGATGAACGAATTGATGTATAGGTGGATGGATGGATGAATAAATGAATGG
+
KKKKKKKKKKKKKKKKKKKKKKKKKKKKKKKKKKKKKKKKKKKKKKKKKKKKKKKKKKKKKKFFKKAKKFFKKKKAKKKKFKFFKKKKKKKKKFKKKK,<KFKK7FKFAAFAFKFA7AF7,AA<,AA<FKF
@ST-E00273:177:HMTTCCCXX:1:1101:11302:23425 BX:Z:AAACACCAGATTCAAG-1
TCCATTCCATGATTATTCCATTGGAATCCATTCAGCGATGATTCCATATGAGTCAATCAATGATGATTCCATATGGGTCCGTTCGATGATAATTCCATTTGATTTCATTCGATGCTTCTATTCTATTC
+
KKKKKKKKKKKKKKKKKKKKKKKKKKFKKKKKKKKKKKKKKKKKKKKKFKKFKKKFKKFKKKKFKKKKKKKKKFKFKKKFFFFKAFKKKKKKKKKKKKKKKKKKKKKKKKKKKKKKKKKKKKFKKFA,<FAA
@ST-E00273:177:HMTTCCCXX:1:1103:30837:64369 BX:Z:AAACACCAGATTCAAG-1
ATACTTATTGTTTTCTCTTAAGTTTATGATTGCCTCTATTTTTGCTTGTTGAGAAAGAAAATGCATGTCTTCTTCATTATGTTCCTCATATCTTCCAGTTTTCCACTCTTTGGTATGCATTCCATTCT
+
KKKKKKKKKKKKK7FKA7FFFK,FFKAFKKKKKKKKKFKFKFKKKAFAFKKKKKKKKKKKKKFKFKKKAKKAFKAKKKKKKFKFKKKKFKFKFKFFFKKFKK,AFFKKKKKKKKFKKKKKK7AFAA,,
@ST-E00273:177:HMTTCCCXX:1:1104:20263:16481 BX:Z:AAACACCAGATTCAAG-1
CATCTGCTCTTCTTTCCTTCTCAGAAAACCTTTTGATGTATGTGCGTGCATGTGCATAGGAGTGTGTGTGCATGAGCATGTACATAGTAGTATATTTTAAAAATTACTGTGAGAACTGTAATTAGCTA
+
KKKKKKKKKKKKKKKKFKKKKKKKKKKKKKKKKKKKKKKFKKKKKKKKKKKKKKKKKKKKKKKKKKAFFKKKKKKKKKKKFKFFFKKKFKKKKKKKKKKKKKKKKKKKKKKFK<FKKKKKKKFKKKKKKKKKKAK
@ST-E00273:177:HMTTCCCXX:1:1106:17330:70188 BX:Z:AAACACCAGATTCAAG-1
TTGTAGGATGCTGAGGTTTGTGCTTCTAATAGTAAGCAGGCTGTCCTTTGTGAATAACAATCACAGATACAGACATCTTCAGACAGTTCAAGGCAAGTCGACTATGATCTCATCAGGGCAGTGGGCTT
+
KKKKKKKKKKKKKKKK,FFFKKKKKKKKK7KKKKKKKKKKKKKKKKFKKKKKKFKKKKKKKKKAKKKKKKKKKKK<AFFKKKKKKKKKAFKKKAFKAFFKFKKKKKKKKKKKKKKKKKFKKK7F,AAAKA
@ST-E00273:177:HMTTCCCXX:1:1113:4208:55965 BX:Z:AAACACCAGATTCAAG-1
GGTCAGGGATGTTACCAGACATCCTATAATGCACAGGAAAACCCTCCCTGACCCACAATGAAACATTGTTAGCCCAAGTAATGCTGAGATTGAGAAACCCTGCTGTAAAGTGTGGATTTTTATCTTGA
+
KKKKKKKKKKKKKKKKKKKKKKKKKKKKKKKKKKKKKKKKKKKKKKKKKKKKKKKKKKKKKKKKKKKKKKKKKKKKKKKKKKKKKKKKKKKKKKKKKKKKKKKKKKKKFKFFFKKKKKKKKKKK<F7
@ST-E00273:177:HMTTCCCXX:1:1206:12205:34201 BX:Z:AAACACCAGATTCAAG-1
TTTAAATTAAGCTCTATGACTCTGTTAATCCTACAAATTTCTTGTTGTTCCCCGTGTCAATTAAGATGTTTTTGCCTGTAAGCCACAGAACATCCAACTAAAAATGAGGGAAATTTATTTCACATAAC
+
KKKKKKKKKKKKKAKKKKKKKKKKKKKKKKKKKKKKKKKKKFKKKFFKKKKKKKFAF<<KKKKKKKKKKFKKKKKKKKKKKKKKKKKKKKKKKKKFKKKKKKKKKKKFFKKKKKKK7<FKKKFKFKK77FA
@ST-E00273:177:HMTTCCCXX:1:1210:5832:12156 BX:Z:AAACACCAGATTCAAG-1
CCACACTATCTCTATCACCACCACTGTCACCAGCATGTCTAGCACCACCACCACCATTACCACCATCATCACCACCGTCATCTCTGTCACACCACCATCACCACCGTGATTGTCACCACCACCATCTC
+
KKKKKKKKKKKKKKKKKKKKKKKKKKKKKKKKKKKKKKKKKKKKKKKKKKKAKFKKKKKKKKKKKKKKKKKKKKKKKKKKKKKKKKKKKKKFFKKKFKKKKKK7AAKKAFKFFFKFFKKKKKKF,FFKFA,<7A
```

**B.**

```
ST-E00273:177:HMTTCCCXX:1:1116:18568:72297      65      chr1    9999    14      14S39M  chr4    190204518       0       TGG
TTCCTCTTCTTGATAACCCTAACCCTAACCCTAACCCTAACCCTAACCCT    AAFFFKKFKKKFKKKKFKKKKKKKKKKKKKKKKAKKKKKKKAFFKFFKKKA7   XF:i:0  PG:Z:M
arkDuplicates     RG:Z:NA12878-1846AF55   XG:f:0.96828    MI:i:49195812   NM:i:0  BX:Z:GTGCGACGTCCCTACT-1
ST-E00273:177:HMTTCCCXX:1:1217:19268:73053      99      chr1    9999    0       103M    =       9999    103     CATAACCCTAA
CCCTAACCCTAACCCTAACCCTAACCCTAACCCTAACCCTAACCCTAACCCTAACCCTAACCCTAACCCTAACCCTAACCCTAACC KKKKKKKKKKKKKKKKKKKKKKKKKKKKKKK
AKKKKKKKKKKKKKKKKKKKKKKKKKKKKKKKKKKKKKKKKKKKKKFFKKK7FKFKKKKFKFFKAFKKFFKAKKKF XA:Z:chr1,+10005,103M,1;       XF:i:1  PG:Z:Mark
Duplicates     RG:Z:NA12878-1846AF55   XG:f:0.0024337  MI:i:52639261   NM:i:1  BX:Z:TAGAAGAAGCCAACAG-1
ST-E00273:177:HMTTCCCXX:1:1210:6979:62312       97      chr1    9999    32      116M    chr5    10005   0       GATAACCCTAA
CCCTAACCCTAACCCTAACCCTAACCCTAACCCTAACCCTAACCCTAACCCTAACCCTAACCCTAACCCTAACCCTAACCCTTACCCGAACCCAACCCC    AAFAFFAKKKKKKK
KKKKFKF,F<FKFFKAFA7FKKKKKFAFAAKFAKA<7FKKFF,,7AFFFKKKFA<F<KKKKFFK7FAKKFAAFAKF,AFAKKKKK,<,<F,<,A,F,7,,7,    XF:i:0  PG:Z:Mark
Duplicates     RG:Z:NA12878-1846AF55   XG:f:0.99938    MI:i:53203580   NM:i:3  BX:Z:TAGGGTTTCTCACATT-1
ST-E00273:177:HMTTCCCXX:1:2224:12347:5089       163     chr1    9999    0       109M42S =       10008   138     CATAACCCTAA
CCCTAACCCTAACCCTAACCCTAACCCTAACCCTAACCCTAACCCTAACCCTAACCCTAACCCCAACCCTAACCCTAACCCCAACCCTAACCATAACAACAACCCAAACCCTAACGA
TGACAACAACCGTCACA AAAFFKKFKKKKKKKKKKKKKKKKKKKKKKKKKKKKKKFKKKKKKKKFKKFKA7FKKFKKKFFFKKFAFF<FAFAA,,7,FF,A,FAFK,AKFA,7<,F<,7,,,,7
,,,<,<,,,,,,,,,7,,,,,,,,,,,,,,,A,,,,,,7,,,,,,, XA:Z:chr5,+11676,2S107M42S,2;   XF:i:1  PG:Z:MarkDuplicates     RG:Z:NA12878
-1846AF55   XG:f:0.47531    MI:i:53366815   NM:i:3  BX:Z:TAGTTGGAGGACGTAC-1
ST-E00273:177:HMTTCCCXX:1:1108:6035:33814       99      chr1    9999    5       13S67M2S        =       9999    67      CGG
ACTGTCGCATGATAACCCTAACCCTAACCCTAACCCTAACCCTAACCCTAACCCTAACCCTAACCCTAACCAG      AAFFFKKKKKKKKKKKKKKKKKKKKKKKKKKKKKKKKKKK
KKKKKKKKKKKKKKKKKKKKKKKKKF,FKKKKKKKKKKKKKKKK,A      XA:Z:chr2,+181275793,15S67M,1;  XF:i:1  PG:Z:MarkDuplicates     RG:Z:NA12
878-1846AF55   XG:f:0.68525    MI:i:56132793   NM:i:0  BX:Z:TCGAAACAGAACGCCA-1
ST-E00273:177:HMTTCCCXX:1:2115:11820:2803       65      chr1    9999    0       14S103M34S      chrX    156030514       0
       ACCTTTACACTCATGATAACCCTAACCCTAACCCTAACCCTAACCCTAACCCTAACCCTAACCCTAACCCTAACCCTAACCCTAACCCTAACCCAAACCCCACCCCTAAC
CACAGCCCACACCCACCCCCCACCCCCCGCCCCAC AAFFFKKKKKKKKKKKKKKKKKKKKKKKKKKAKKKKKKKFKKAKKKKKFFFKF,A<FKK7A7,FA77FKAK77FFKF<F,,FFA7<F7
<7,,FF7,F,F7F,7<FAK,7(((,,7,,A,,,,7,<,(,,7,,,((,<F7,,A,(A((,,(,( XA:Z:chr5,+11694,16S91M2D10M34S,3;      XF:i:0  PG:Z:MarkD
uplicates     RG:Z:NA12878-1846AF55   XG:f:0.64452    MI:i:59118095   NM:i:3  BX:Z:TGAGCATAGACGCCCT-1
ST-E00273:177:HMTTCCCXX:1:2202:6116:58075       1097    chr1    9999    0       13S86M52S       *       0       0       CGG
AAAGTCTTCTGATAACCCTAACCCTAACCCTAACCCTAACCCTAACCCTAACCCTAACCCTAACCCTAACCCTAACCCTAACCCTAACCCCCAACCCCACCCCCCCAACCCTCACC
CCACCCCTCACCCTAACCCCACCCC AAFFFKKKKKKKKKKKKKKKKKKKKKKKKKKKKKKKKKKKKKKKKKKKKKKKKKKKKKKKKKKAKKKKKAFFKKK,<FAKK,7FFKK,AFFKKFA<F
FAA<((,A7,7,(((,7<(,,7,,,,,7,A(,(<<(,(,,<A,,,7<<,,(,(, XA:Z:chr5,+11700,15S85M51S,0;   XF:i:1  PG:Z:MarkDuplicates     RG:Z
:NA12878-1846AF55   XG:f:0.65534    MI:i:60645204   NM:i:0  BX:Z:TGCTATTAGGTCTGTT-1
ST-E00273:177:HMTTCCCXX:1:2104:12581:10240      99      chr1    9999    40      13S43M  =       9999    43      TTCTCAACAGT
CTGATAACCCTAACCCTAACCCTAACCCTAACCCTAACCCTAACC        AAFFFKKKKKKKKKKKKKKKKKKKKKKKKKKKKKKKKKKKKKKKKKKKKKKKKKK        XF:i:1
  PG:Z:MarkDuplicates     RG:Z:NA12878-1846AF55   XG:f:1  MI:i:66357267   NM:i:0  BX:Z:TTGTTGTAGCTATCTG-1
ST-E00273:177:HMTTCCCXX:1:1202:30797:32092      97      chr1    9999    5       14S70M  chr12   10195   0       TGGTTCCTCTT
CTTGATAACCCTAACCCTAACCCTAACCCTAACCCTAACCCTAACCCTAACCCTAACCCTAACCCTA    AAFFFKKKKKKKKKKKKKKKKKKKKKKKKKKKKKKKKKKKKKKKK
KKKKKKKKKKKKKKKKKKKKKKKKKKKKKKKKKKKKKKKK    XA:Z:chr1,+10037,16S68M,0;      XF:i:1  PG:Z:MarkDuplicates     RG:Z:NA12878-1846
AF55   XG:f:0.69223    MI:i:67068505   NM:i:0  BX:Z:TTTGCTAAGGAGTGTC-1
ST-E00273:177:HMTTCCCXX:1:2101:11403:60519      147     chr1    9999    11      13S49M  =       9999    -49     GACTGTAAGGT
AGGATAACCCTAACCCTAACCCTAACCCTAACCCTAACCCTATCCCTAACC  7KKKFFFA7A77FFAFKKKKKFKKKKKKFA<A<A,FKAKKKFKKKKF,AF7KF,KAKFAFAA  XF:i:1
  PG:Z:MarkDuplicates     RG:Z:NA12878-1846AF55   XG:f:0.97709    MI:i:39084628   NM:i:1  BX:Z:GAGTCCGAGAGCGAAA-1
ST-E00273:177:HMTTCCCXX:1:1217:19268:73053      147     chr1    9999    0       103M    =       9999    -103    CATAACCCTAA
CCCTAACCCTAACCCTAACCCTAACCCTAACCCTAACCCTAACCCTAACCCTAACCCTAACCCTAACCCTAACCCTAACCCTAACC A,,<AKF7F,AKKFKA7KKFAF<KFFFFAK
KKKKFKKKKKFKKFKAFKKKKAKKKKKKKKKKKKKKKKKKKKKKKKKKKKKKKKAKKKKFKKKKKKFKKKFFAAA XA:Z:chr1,-10005,103M,1;       XF:i:1  PG:Z:Mark
Duplicates     RG:Z:NA12878-1846AF55   XG:f:0.0024366  MI:i:52639261   NM:i:1  BX:Z:TAGAAGAAGCCAACAG-1
ST-E00273:177:HMTTCCCXX:1:1108:6035:33814       147     chr1    9999    5       13S67M2S        =       9999    -67     CGG
ACTGTCGCATGATAACCCTAACCCTAACCCTAACCCTAACCCTAACCCTAACCCTAACCCTAACCCTAACCAG      A,,KKAKKK<FK<A7KAFKKKKFAKKKKKFKKKKKKKK
FKF<KKKKFAKKKKFFKKKKKFKKFFKKKKKKKKKKKKFFFA,A      XA:Z:chr2,-181275793,15S67M,1;  XF:i:1  PG:Z:MarkDuplicates     RG:Z:NA12
878-1846AF55   XG:f:0.68398    MI:i:56132793   NM:i:0  BX:Z:TCGAAACAGAACGCCA-1
```

**Supplemental Figure 2. (A)**. Illustration of the unified FASTQ format. Each read contains the barcode field “BX:Z:barcode” following the read id. The length of barcode sequence are 16 bp, 30bp and 18bp for 10x, stLFR and TELL-Seq, respectively. **(B)**. Illustration of the barcode information in the alignment file. The barcode information is presented using the “BX:Z:barcode” format in the optional filed of alignment files.
