## Supplementary figures and images for "LRTK: A platform agnostic toolkit for linked-read analysis of both human genomes and metagenomes"

### SupplementaryFigure2.pdf

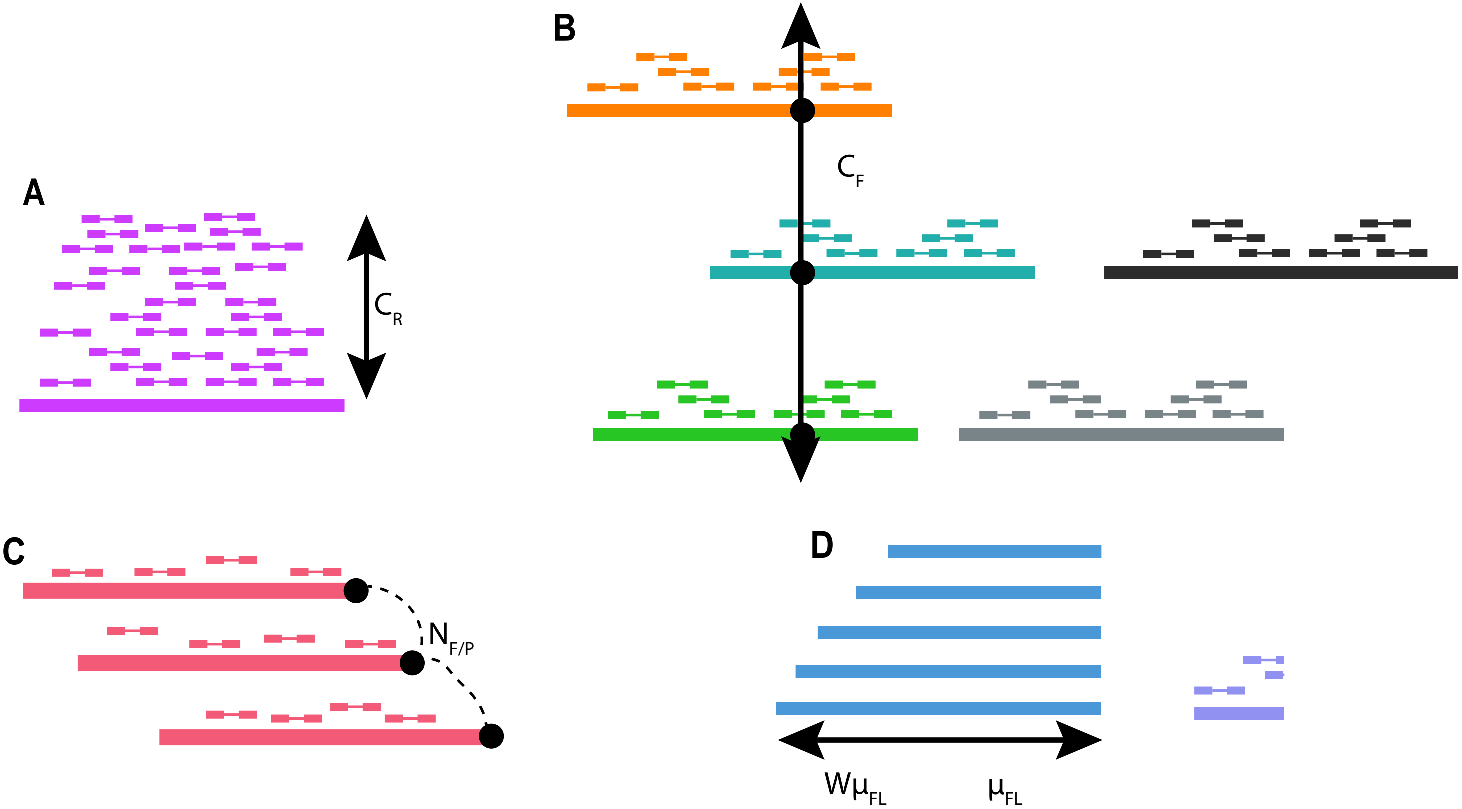

### SupplementaryFigure3.pdf

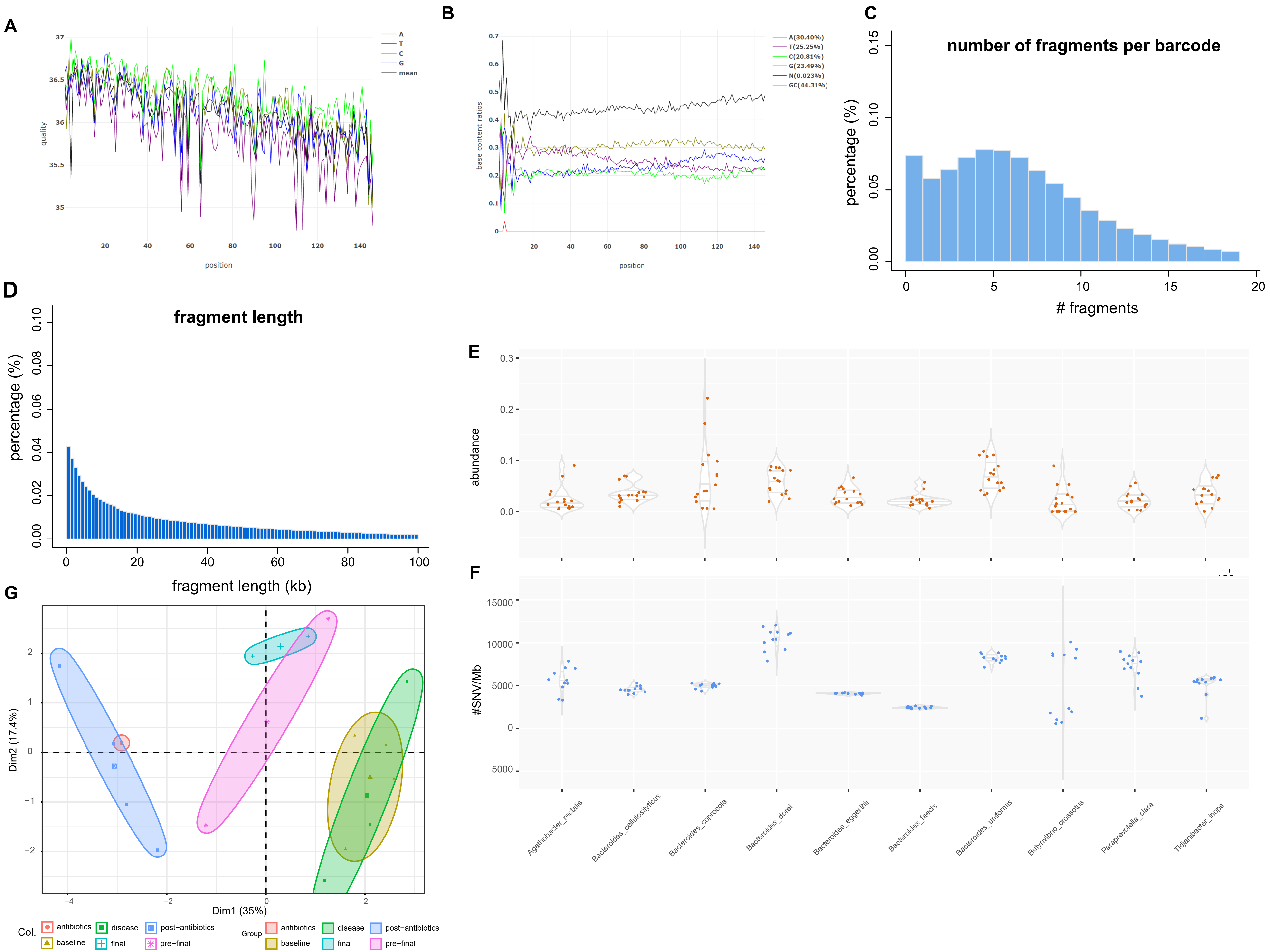

### SupplementaryFigure4.pdf

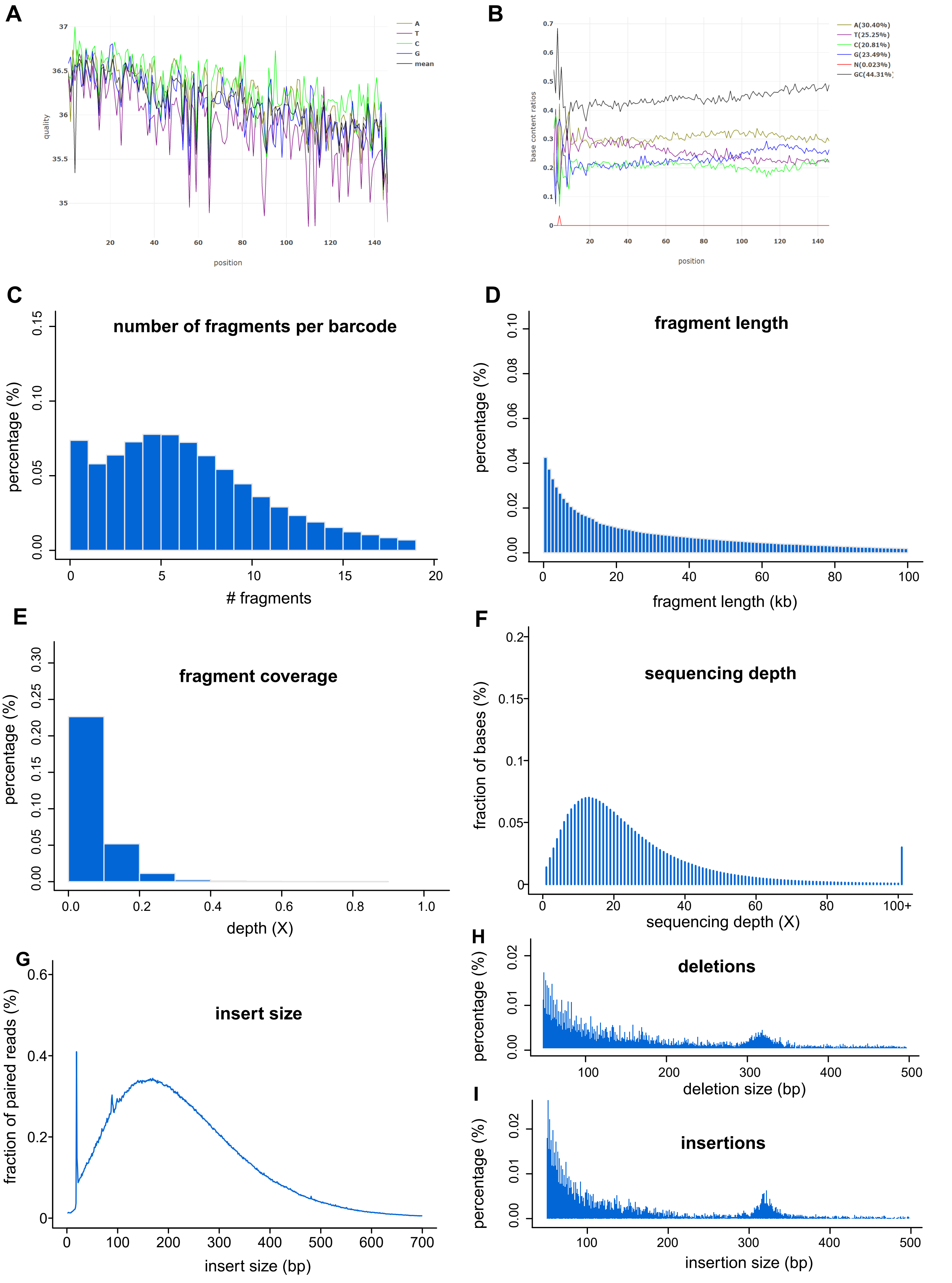
